# Targeted attenuation of elevated histone marks at *SNCA* alleviates α-synuclein in Parkinson’s disease

**DOI:** 10.1101/2020.02.13.947465

**Authors:** Subhrangshu Guhathakurta, Jinil Kim, Sambuddha Basu, Evan Adler, Goun Je, Mariana Bernardo Fiadeiro, Yoon-Seong Kim

**Author notes:** Corresponding author: Yoon-Seong Kim, M.D., Ph.D., Associate Professor, Division of Neurosciences, Burnett School of Biomedical Sciences, College of Medicine, University of Central Florida, 6900 Lake Nona Blvd., Orlando, FL 32827.

## Abstract

Epigenetic de-regulation of α-synuclein plays a key role in Parkinson’s disease (PD). Analysis of the *SNCA* promoter using the ENCODE database revealed the presence of important histone posttranslational modifications (PTMs) including transcription-promoting marks, H3K4me3 and H3K27ac, and repressive mark, H3K27me3. We investigated these histone marks in postmortem brains of controls and PD patients and observed that only H3K4me3 was significantly elevated at the *SNCA* promoter of the substantia nigra (SN) of PD patients both in punch biopsy as well as in NeuN positive neuronal nuclei samples. To understand the importance of H3K4me3 in regulation of α-synuclein, we developed CRISPR/dCas9-based locus-specific H3K4me3 demethylating system where the catalytic domain of JARID1A was recruited to the *SNCA* promoter. This CRISPR/dCas9 SunTag-JARID1A significantly reduced H3K4me3 at *SNCA* promoter and concomitantly decreased α-synuclein both in the neuronal cell line SH-SY5Y and idiopathic PD-iPSC derived dopaminergic neurons. In sum, this study indicates that α-synuclein expression in PD is controlled by *SNCA’s* histone PTMs and modulation of the histone landscape of *SNCA* can reduce α-synuclein expression.

## Introduction

Parkinson’s disease (PD) is the second most prevalent neurodegenerative disease affecting nearly one million people worldwide. In the USA alone, 60,000 patients are diagnosed with PD each year (Parkinson’s Foundation). PD is a late-onset disease that destroys 70 to 80 percent of dopaminergic neurons in the susbstantia nigra pars compacta region of the midbrain before motor symptoms are typically noticeable (Heisters, 2011; Morrish et al, 1998; Postuma et al, 2010). The disease is typically idiopathic, but cases with known genetic components account for around 10 percent of reported cases (Gasser, 2009). Of note, α-synuclein is one of the primary proteins linked with PD, and it has been identified as playing a role in both genetic and non-genetic cases.

The first evidence of α-synuclein involvement in PD came from several familial studies that showed individuals harboring coding region mutations in *SNCA,* the gene encoding α-synuclein, including A53T, E46K, and A30P, had early-onset disease with an autosomal dominant pattern of inheritance (Kruger et al, 1999; Polymeropoulos et al, 1997; Zarranz et al, 2004). Later, more familial *SNCA* mutations were identified including H50Q, G51D, A18T, and A29S, all of which have been shown to affect disease manifestation and aggregation of α-synuclein (Appel-Cresswell et al, 2013; Hoffman-Zacharska et al, 2013; Lesage et al, 2013). Moreover, familial PD cases with locus multiplications, such as duplication and triplication of *SNCA,* exhibited severe forms of the disease and early onset (Chartier-Harlin et al, 2004; Singleton et al, 2003). Duplication or triplication of *SNCA* could produce higher amounts of α-synuclein in neurons, which might account for the aggressive form of the disease observed in those patients (Fuchs et al, 2008; Miller et al, 2004).

The central role of α-synuclein in the pathogenesis of idiopathic PD was supported by the discovery that aggregates of α-synuclein are major components of Lewy Bodies, the protenaceous intracytoplasmic inclusions in dopaminergic neurons found in postmortem brains of PD patients (Spillantini et al, 1997). In addition, a single-cell study using laser capture microdissection found significantly higher levels of α-synuclein transcripts in PD postmortem brains (Grundemann et al, 2008). Together, these studies indicate regulation of the *SNCA* gene is important and elevated levels of α-synuclein could be a key factor in development of PD and other synucleinopathies.

Numerous studies have investigated the make-up, formation, and propagation of α-synuclein aggregates in PD (Giraldez-Perez et al, 2014; Stefanis, 2012); however, little is known about the mechanisms underlying transcriptional or epigenetic de-regulation of *SNCA* in PD. Except for two conflicting reports that investigated hypomethylation of the intron 1 CpG island of *SNCA* in PD and control postmortem brain samples, studies of epigenetic factors regulating the α-synuclein gene in PD have been limited (Guhathakurta et al, 2017a; Guhathakurta et al, 2017b).

The ENCODE database now allows for in-depth analysis of the epigenetic environment of genes (Consortium, 2012). And while the importance of histone posttranslational modifications (PTMs) in regulation of gene expression has been established, to date, no studies have comprehensively investigated the potential role of histone PTMs in regulation of *SNCA* in PD (Guhathakurta et al, 2017a).

In this study, we investigated the epigenetic environment of *SNCA* with special emphasis on histone PTMs that are potentially enriched in the regulatory region of the gene. Interestingly, we observed a significant increase in one histone PTM, histone H3 lysine 4 trimethylation (H3K4me3) at the *SNCA* promoter in postmortem PD samples. This transcription-initiating histone mark is one of the major determinants of gene transcription (Barski et al, 2007). Reduction of H3K4me3 in neuronal cell lines and PD-derived induced pluripotent stem cell lines (iPSCs) using the dead Cas9-Suntag system-mediated locus-specific approach decreased levels of α-synuclein. Results of this epigenetic investigation could open new avenues for therapeutic intervention to reduce α-synuclein expression by preventing enrichment of H3K4me3 in the *SNCA* gene.

## Results

### Analysis of histone architecture of human *SNCA* in postmortem midbrain samples

Human *SNCA* consists of six coding exons and two upstream non-coding exons and spans a 114-kb region in chromosome 4 (Guhathakurta et al, 2017a). We first searched *in silico* for any histone marks in the adult human brain substantia nigra (SN) region using the Roadmap Epigenomics Database (http://www.roadmapepigenomics.org/data/tables/adult). In total, the database lists seven histone PTMs in the SN region of the brain from two adult postmortem samples: H3 lysine 27 trimethylation (H3K27me3), histone H3 lysine 36 trimethylation (H3K36me3), histone H3 lysine 4 monomethylation (H3K4me1), histone H3 lysine 4 trimethylation (H3K4me3), histone H3 lysine 9 trimethylation (H3K9me3), histone H3 lysine 9 acetylation (H3K9ac), and histone H3 lysine 27 acetylation (H3K27ac). However, for *SNCA,* not all seven histone PTMs were enriched in their cohort of tissues analyzed. H3K36me3, the histone mark associated with full-length transcription, was enriched throughout the gene body, ensuring *SNCA* is actively transcribed in the SN region. Most interestingly, we observed that H3K4me3, H3K27ac, and H3K27me3 were the three histone PTMs preferentially enriched in the primary regulatory regions of *SNCA—*the area ranging from approximately -1 kb to +1.5 kb of the transcription start site (TSS). This transcriptionally important region includes the promoter, its upstream regions, and the intron 1 area of the gene (chr4:90,757,022-90,759,612 bp; GRCh37/hg19). Fig 1A (and Fig EV1) shows the distribution pattern of the three histone marks around the promoter region for exon 2, the first coding exon. Both H3K4me3 and H3K27ac are transcription favoring histone marks. H3K4me3 is the principal mark associated with transcription initiation, and H3K27ac is an active enhancer-associated mark often enriched at the promoter. By contrast, H3K27me3 is associated with gene repression.

**Figure 1.**
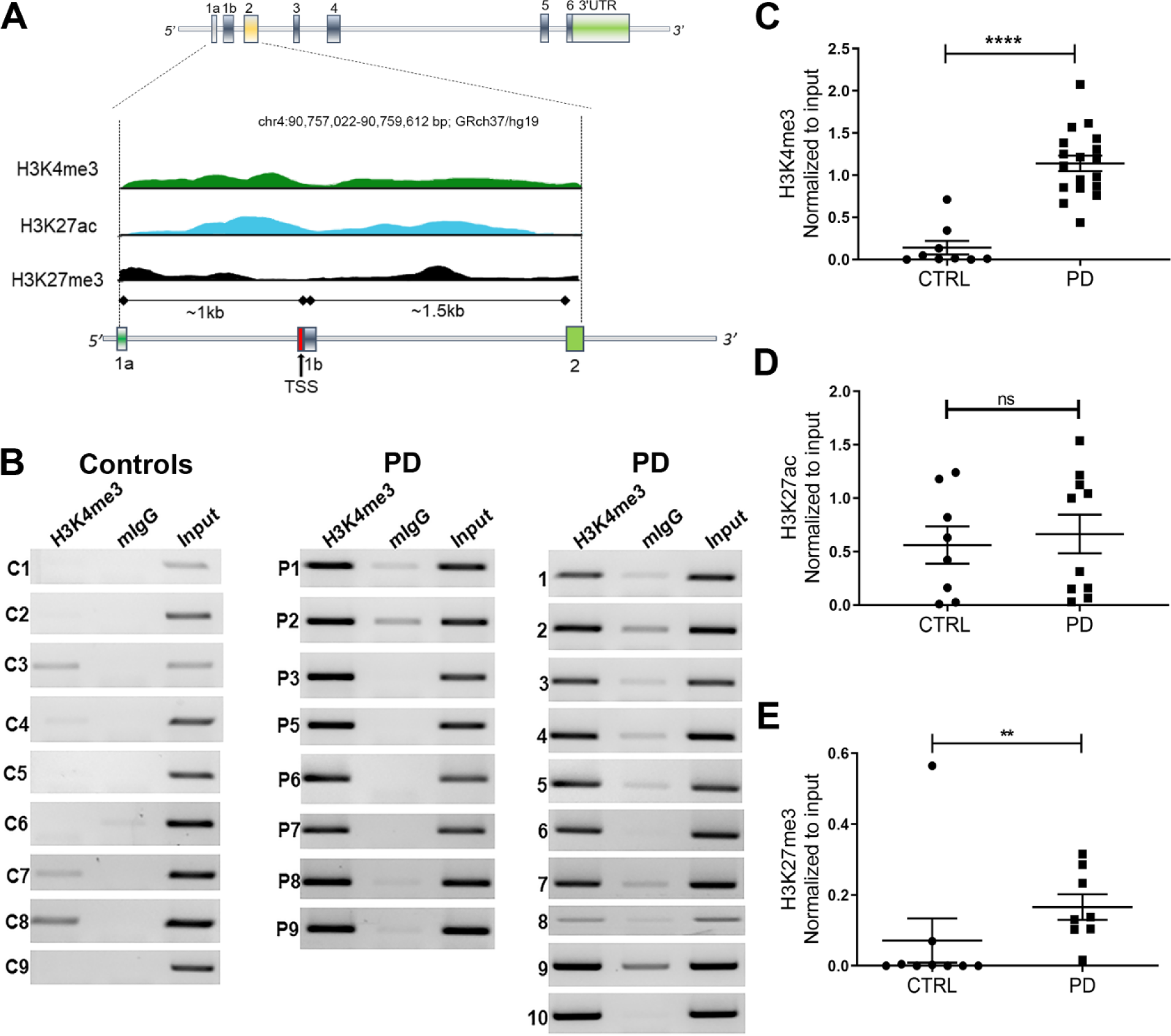
Parkinson’s disease patients harbor high levels of H3K4me3 at the α-synuclein promoter. **A** Human *SNCA* gene contains six coding exons and two 5’ non-coding exons. The exons are represented by vertical colored boxes. The region from exon 1a to exon 2 (∼2.5 kb) is scaled up to show the distribution of histone PTMs. Distribution of the histone PTMs from the SN region of one donor brain sample is shown. Peaks of three different histone PTMs, H3K4me3 (green), H3K27ac (blue), and H3K27me3 (black), at the regulatory region of *SNCA* were adopted from Roadmap Epigenomics Database. For a detailed view, please see Fig EV1 where screenshot of the original figure is shown. The TSS is indicated by a red vertical bar and distances of exon 1a and 1b from the TSS are indicated. **B** ChIP gel images showing the relative enrichment by H3K4me3 in controls (n=9) and PD patients (n= 18). PCR amplified a 188-bp region of *SNCA* from intron 1 where H3K4me3 peak was at its optimum. Mouse IgG (mIgG) was used as control and the bands were normalized by unbiased amplification from respective inputs. **C** Relative intensities calculated from the gel images. Graph shows that H3K4me3 was significantly enriched at the upstream regulatory region of *SNCA* in PD compared to control subjects. **D** Difference in enrichment of H3K27ac in PD (n= 10) and controls (n=9) was not significant for the same region of the gene. **E** The transcriptional repressive histone mark, H3K27me3, was significant in PD samples compared to controls. *\*P* < 0.05, \*\**P* < 0.01, \*\*\**P* < 0.001, and \*\*\*\**P* < 0.0001, ns = non-significant. Data were analyzed using non-parametric t-test followed by Mann-Whitney post-hoc corrections. Two-tailed p-values were calculated for all. Data information: Data represent mean ± standard error of the mean.

In *SNCA*, we observed both the transcription-promoting marks, H3K4me3 and H3K27ac, had sharp peaks around the TSS and surrounding areas while H3K27me3 had an overall low-level distribution at the promoter and intron 1 areas. Therefore, we next analyzed these three histone marks in our cohort of postmortem midbrain tissue samples of PD and matched controls specifically from the SN region using chromatin immunoprecipitation (ChIP) (Fig 1A-E; Fig EV1). We found that H3K4me3 was significantly enriched at the *SNCA* regulatory region in PD samples compared to controls (p<0.0004) (Fig 1B-C). All the PCRs were performed for 35 cycles to keep the PCR products in unsaturated condition. On the other hand, no significant difference was observed for H3K27ac between control and PD subjects (Fig 1D, Fig EV2A). We also found relatively higher enrichment of H3K27me3 in PD; however, the difference in H3K27me3 between control and PD was less prominent and significant than the difference in H3K4me3 between the two groups (Fig 1E; Fig EV2B). Enhancer activity of the *SNCA* intron 4 region has been reported (Guhathakurta et al, 2017a; Soldner et al, 2016). As mentioned above, enhancer regions are often enriched by H3K27ac as shown in the Roadmap epigenetic data for *SNCA* (Guhathakurta et al, 2017a; Soldner et al, 2016). However, we did not find any significant difference in the enrichment of H3K27ac in the intron 4 enhancer region between control and PD subjects (Fig EV3).

### Relative enrichment of H3K4me3 in the neuronal population of postmortem PD brain

Cellular heterogeneity in the brain is a major obstacle when studying neuronal epigenetic architecture. We further investigated if the observed difference in H3K4me3 between control and PD is maintained in neuronal populations in the SN region. We isolated 20,000 neuronal nuclei from the SN region of a subset of brain samples (6 control and 7 PD) using NeuN-based fluorescence-activated nuclei sorting (FANS). NeuN antibody is a widely used neuron-specific antibody that preferentially binds to Fox3 transcription factor in the neuronal nuclei (Kim et al, 2009; Mullen et al, 1992). We were also able to amplify neuron-specific transcripts (NeuN, synaptophysin, and α-synuclein) and GFAP for astrocytes from the isolated NeuN+ nuclei, confirming the purity of neuron-specific isolation (Fig 2A-B). As expected, neuronal nuclei isolated from equal amounts of tissue from the SN region yielded much higher numbers of NeuN+ nuclei in controls compared to PD (Fig 2). Using the same primer set used for whole-tissue ChIP of H3K4me3 at the *SNCA* promoter/intron 1 region, we found that NeuN+ nuclei from PD samples demonstrated significantly higher enrichment of H3K4me3 compared to controls (p =0.015) (Fig 2C-D).

**Figure 2.**
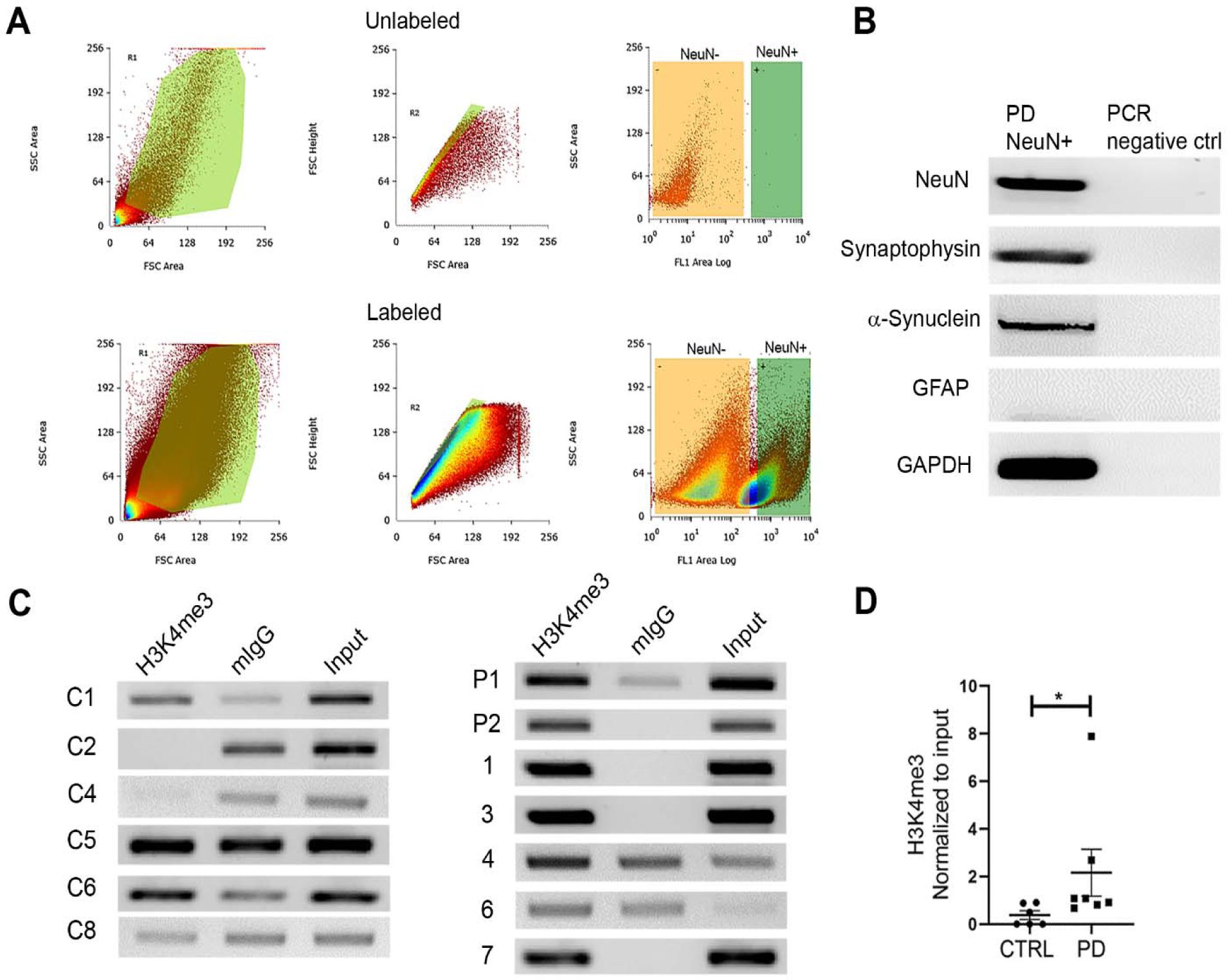
NeuN-positive neurons from the SN demonstrate high enrichment by H3K4me3. **A** 20,000 NeuN labeled neuronal nuclei were collected by fluorescent activated nuclei sorting (FANS) from control (n = 6) and PD (n= 7) SN tissues. The NeuN-positive nuclei were labeled by anti-rabbit IgG secondary antibody tagged with Alexa fluor 488. The top and bottom panels show representative sort gating windows from an unlabeled and labeled patient sample, respectively. The left column represents size versus granularity gatings. The samples were then gated for singularity using forward scatter height (Y-axis) versus forward scatter area (X-axis), and lastly, singularly gated nuclei were sorted based on NeuN positivity (X-axis; FL1 channel) versus side scatter (Y-axis). The green rectangular quadrants on the right column of both panels represent the NeuN+ region, as unstained sample did not show any significant representation. Therefore, from each of the stained samples, 20,000 bright NeuN+ nuclei (green rectangle gate) were sorted and collected. **B** Gel images from RT-PCR show purity of the sorted nuclei from a representative PD sample. Nuclear RNA was isolated, cDNA was generated and pre-amplified (see Materials and Methods for details) before target-specific PCR. Neuron specific genes (NeuN, synaptophysin) and astrocyte specific gene (GFAP), α-synuclein, and GAPDH were amplified from the isolated nuclei. **C** ChIP was performed on the 20,000 isolated nuclei against H3K4me3 from all samples. Gel images represent the ChIP-based PCR amplification. The same primer pair was used to amplify the target region on *SNCA* as Fig EV1. Mouse IgG was used as a control. **D** The graph represents the statistically significant difference of relative H3K4me3 enrichment between PD and controls. Neuronal nuclei from PD brain samples show significantly higher enrichment of H3K4me3 at *SNCA* intron 1 (p=0.01) compared to controls. *\*P* < 0.05. Data were analyzed using non-parametric t-test followed by Mann-Whitney post-hoc corrections. Two-tailed p-values were calculated for all. Data information: Data represent mean ± standard error of the mean.

### Correlation between H3K4me3 levels and high levels of α-synuclein in PD patients

We evaluated α-synuclein levels in midbrain SN tissues from control and PD subjects by western blot analysis. The entire cohort of samples used for analyzing enrichment of H3K4me3 was used for α-synuclein expression. We found that α-synuclein was significantly higher in PD compared to controls (p<0.05) (Fig 3A-B). We then compared whether α-synuclein levels could be correlated with corresponding H3K4me3 enrichment. Toward this end, we calculated the median level of α-synuclein among all study subjects (0.43) and found that 9 PD subjects and 3 controls had higher α-synuclein levels than the median. Interestingly, higher levels of α-synuclein in those subjects were significantly correlated with corresponding H3K4me3 enrichment at the gene promoter (Spearman correlation coefficient (R), 0.71, p =0.01) (Fig 3C). This further established that enrichment of H3K4me3 at the *SNCA* promoter could account for higher expression of α-synuclein.

**Figure 3.**
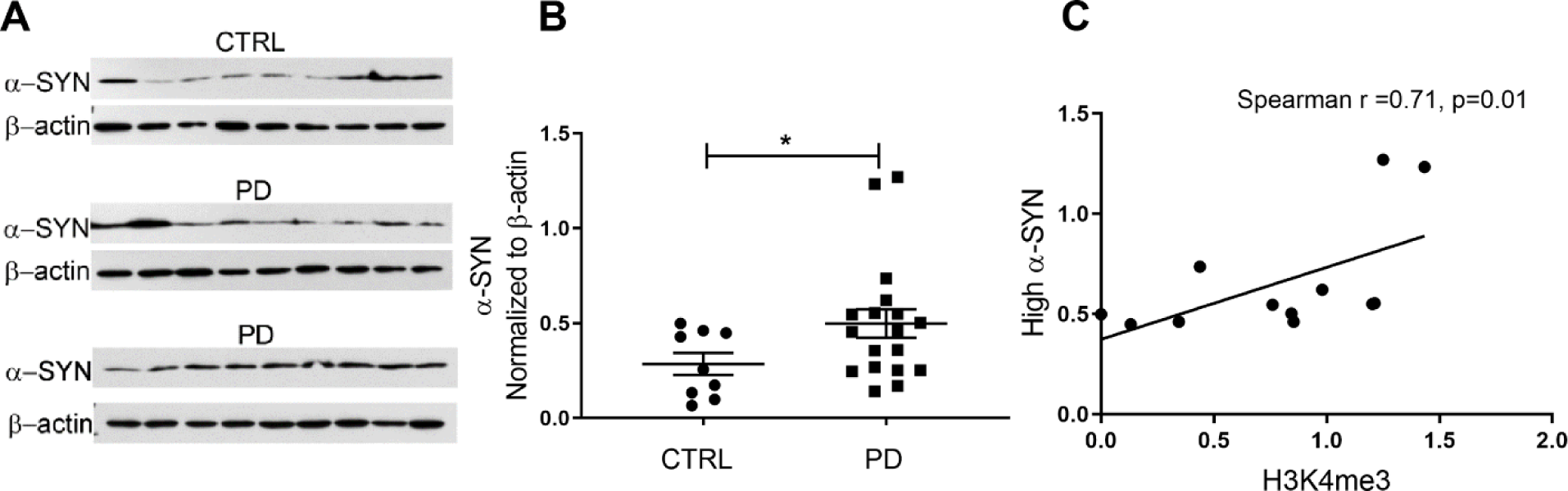
PD patients exhibit high expression of α-synuclein. **A** Western blot gel images showing α-synuclein levels in SN tissues from control (n= 9) and PD subjects (n = 18). **B** The relative levels of α-synuclein between the two groups were evaluated statistically. PD subjects had significantly higher levels of α-synuclein compared to controls. **C** The normalized levels of α-synuclein in the entire cohort were divided into high (n = 12) and low levels. The cut off value for determining the threshold was set at the median from all the subjects. The high levels of α-synuclein were plotted with corresponding H3K4me3 values of those subjects. There was a significant correlation between high levels of α-synuclein and H3K4me3. \**P* < 0.05, \*\**P* < 0.01, ns = non-significant. Data in (**B**) were analyzed using non-parametric t-test followed by Mann-Whitney post-hoc corrections and two-tailed p-values were calculated. Spearman rank correlation test was performed in (**C**). Data information: Data represent mean ± standard error of the mean.

### Design of CRISPR/dCas9 SunTag-JARIDIA system and its recruitment at the *SNCA* promoter

Because the histone mark H3K4me3 was significantly enriched at the *SNCA* promoter in PD patients and was correlated with high levels of α-synuclein in all study subjects, we aimed to investigate whether removal of H3K4me3 at the *SNCA* promoter affected α-synuclein levels. To this end, we searched for an epigenetic eraser that could efficiently remove trimethylation at lysine 4 from the histone H3 tail. Histone lysine demethylases (KDM) are specific “erasers” for methylated lysine residues on histones (Hyun et al, 2017). KDM5A, also known as JARID1A (Jumonji, AT-rich interactive domain, member 1A), has been shown to have specific demethylating activity at H3K4me3 (Horton et al, 2016; Klose et al, 2007). JARID1A contains several conserved domains including n- and c-terminal JmjN and JmjC domains, a DNA binding ARID domain, three PHD domains, a Zn^2^+ binding domain, and a PLU domain (Horton et al, 2016). The first 797 amino acids containing JmjN, ARID, PHD1, JmjC, and Zn^2^+ domains were identified as catalytic because overexpression of a recombinant construct containing these domains was sufficient to demethylate H3K4me3 *in vitro* (Horton et al, 2016). Based on this information, we selected JARID1A as the H3K4me3-demethylase for our next set of experiments. To recruit a JARID1A catalytic domain to the *SNCA* promoter, we used the CRISPR/dCas9 technology-based SunTag or SuperNova system by replacing the synthetic transcription activator, VP64, with the JARID1A catalytic domain (Fig 4A). Originally, the SunTag system was designed to activate gene expression (up to 300X) by recruiting multiple copies (10 or 22) of VP64 tagged with a single chain variable fragment (scFV) that specifically binds to a GCN4 peptide at the target gene promoter (Tanenbaum et al, 2014). Upon co-expression of a dead-Cas9 (dCas9) containing c-terminal repeats of 10 or 22 GCN4 peptides (10x or 22x GCN4) together with genomic locus-specific small guide RNAs (sgRNAs), target gene-specific accumulation of VP64 is achieved, ensuring robust transcriptional activation (Tanenbaum et al, 2014). We sub-cloned the scFV-sfGFP-JARID1A catalytic domain into a lentiviral system where the transgene is expressed under the CMV immediate enhancer element (Fig 4B). The entire sequence of this novel vector system is provided in Fig EV10. Morita *et al*. previously showed that to avoid steric hindrance between adjacent effector molecules and for proper recruitment of larger effectors such as TET1 catalytic domain, a minimum 22 amino acid spacer is necessary between each GCN4 unit (Morita et al, 2016). Therefore, we also used this dCas9-5xGCN4 system, which could recruit five JARID1A molecules at the *SNCA* promoter site when co-overexpressed with scFV-sfGFP-JARID1A and sgRNA (Fig 4A). We designed nine guide RNAs upstream of the TSS of *SNCA* where the peaks of H3K4me3 were relatively higher (Fig 4C). Guide RNA design was carried out using Broad Institute’s genomic perturbation platform to ensure selection of only those sequences that had 0 to 1 unintended potential off-targets elsewhere in the genome (see Materials and Methods for details). The list of sgRNAs with sequences and their relative locations are provided in Table EV3.

**Figure 4.**
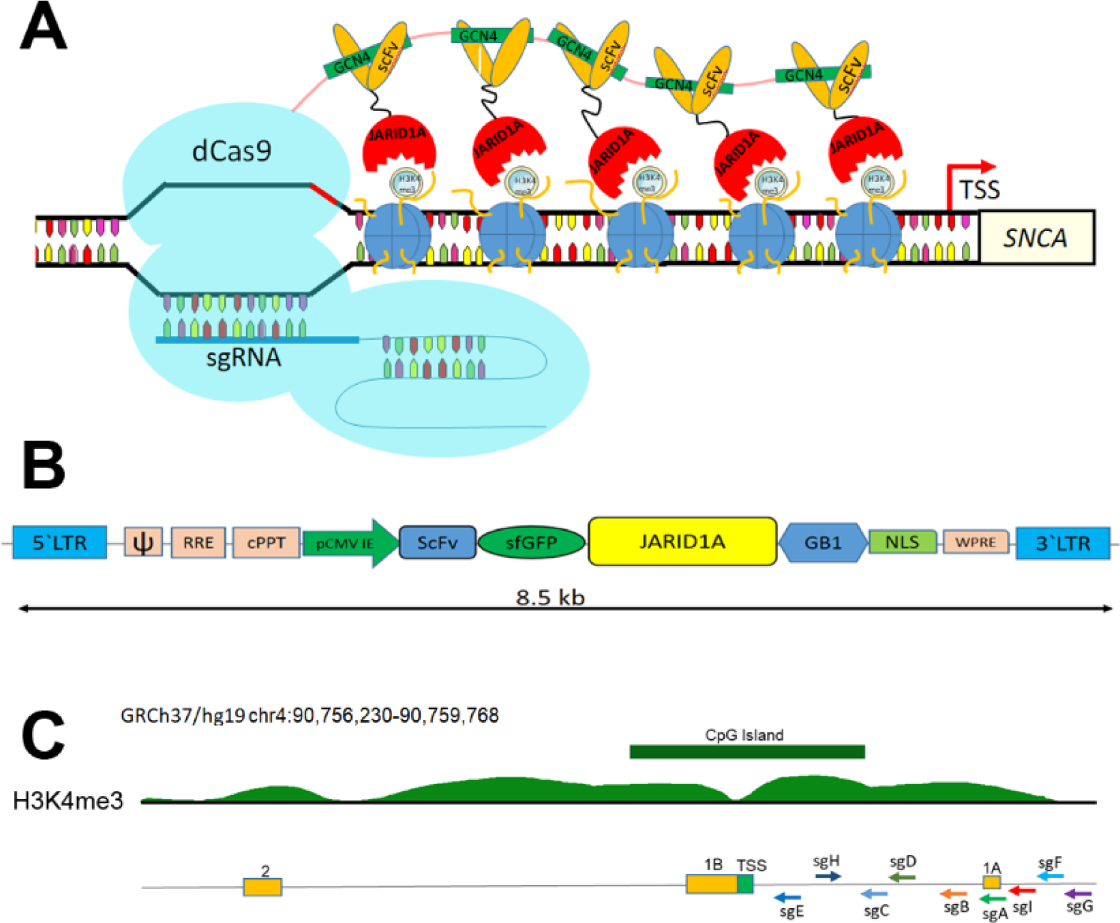
Design of CRISPR/dCas9 based SunTag-JARIDIA system. **A** Schematic diagram shows how the SunTag-JARID1A system is recruited at the *SNCA* promoter. The dCas9-5xGCN4, scFV-JARID1A, and sgRNA plasmids are co-overexpressed in the cells. The dCas9-5xGCN4 is recruited to the *SNCA* promoter as directed by the specific sgRNA. Five scFV-JARID1A molecules in turn recognize the GCN4 polypeptide sequences of dCas9. Upon recruitment of the entire system, SunTag-JARID1A demethylates H3K4me3 at the target region. **B** Structure of pLvx-scFV-sfGFP-JARID1A. The scFV-sfGFP-catalytic domain of JARID1A was sub-cloned into a lentiviral vector. The distance between the 5’ and 3’ Long Terminal Repeats in the vector is 8.5 kb. The components of the plasmids are as follows: V, packaging signal; RRE, rev response element; cPPT, central polypurine tract; pCMV IE, immediate early cytomegalovirus promoter; scFV, single chain variable fragment; sfGFP, super folder GFP; JARID1A; catalytic domain of JARID1A; GB1, solubility tag protein; NLS, nuclear localization signal; WPRE, woodchuck hepatitis virus (WHP) posttranscriptional regulatory element. **C** Relative positions of the nine sgRNAs. A ray diagram exhibits the locations of the nine sgRNAs in relation to the upstream regulatory regions of *SNCA* and the H3K4me3 and H3K27me3 peaks. The locations of the sgRNAs are as follows with respect to the TSS: sgA, 1,117 bp; sgB, 836 bp; sgC, 747 bp; sgD, 700 bp; sgE, 153 bp; sgF, 1,454 bp; sgG, 1,537 bp; sgH, 132 bp; sgI, 1,320 bp. The two non-coding exons (1A and 1B) along with the first coding exon (exon 2) are also shown. GRCh37/hg19 contig was used to describe the location of the gene and associated distribution of histone PTMs. The histone peaks shown were from midbrain regions of two adult postmortem human samples.

To determine if dCas9-5xGCN4 and scFV-sfGFP-JARID1A could form a complex when expressed together, we transiently co-overexpressed these constructs in HEK293 cells and immunoprecipitated using anti-Cas9 antibody and further blotted against anti-GFP antibody (Fig EV4). We observed that cells expressing both dCas9-5xGCN4 and scFV-sfGFP-JARID1A were successfully immunoblotted against anti-GFP antibody followed by Cas9-mediated precipitation, demonstrating that scFV-sfGFP-JARID1 A forms a stable complex with dCas9-5xGCN4 when co-expressed (Fig EV4A-B). By contrast, the cells expressing only dCas9-5xGCN4 could not pull down anything else, confirming specificity.

### Significant reduction in α-synuclein expression by removal of H3K4me3 at the *SNCA* promoter

SH-SY5Y cells, a neuronal cell line, exhibit a relatively high level of α-synuclein expression with higher occupancy of H3K4me3 at the *SNCA* promoter (Fig 5B). Therefore, we chose this cell line to see if recruitment of JARID1A at the promoter region of *SNCA* could reduce its expression. First, we created stable SH-SY5Y cell lines expressing dCas9-5xGCN4, which was confirmed by PCR using primers spanning the GCN4 regions (Fig EV5). Using these cells, we further developed cell lines that stably express each sgRNA to recruit the SunTag system precisely at the targeted genomic locations. All nine sgRNA cell lines were able to recruit dCas9-5xGCN4 precisely at distinct locations of the *SNCA* promoter (Fig EV6). Finally, we stably transfected scFV-sfGFP-JARID1A in these pre-selected cell lines which were already expressing dCas9-5xGCN4 and individual sgRNAs (see Materials and Methods for details). Out of nine different sgRNA-cell lines, only two sgRNAs, sgA and sgD, successfully reduced H3K4me3 from the *SNCA* promoter (Fig EV7). Between sgA and sgD, sgA was selected for the rest of the study, as it completed removed H3K4me3 from the *SNCA* promoter of SH-SY5Y.

**Figure 5.**
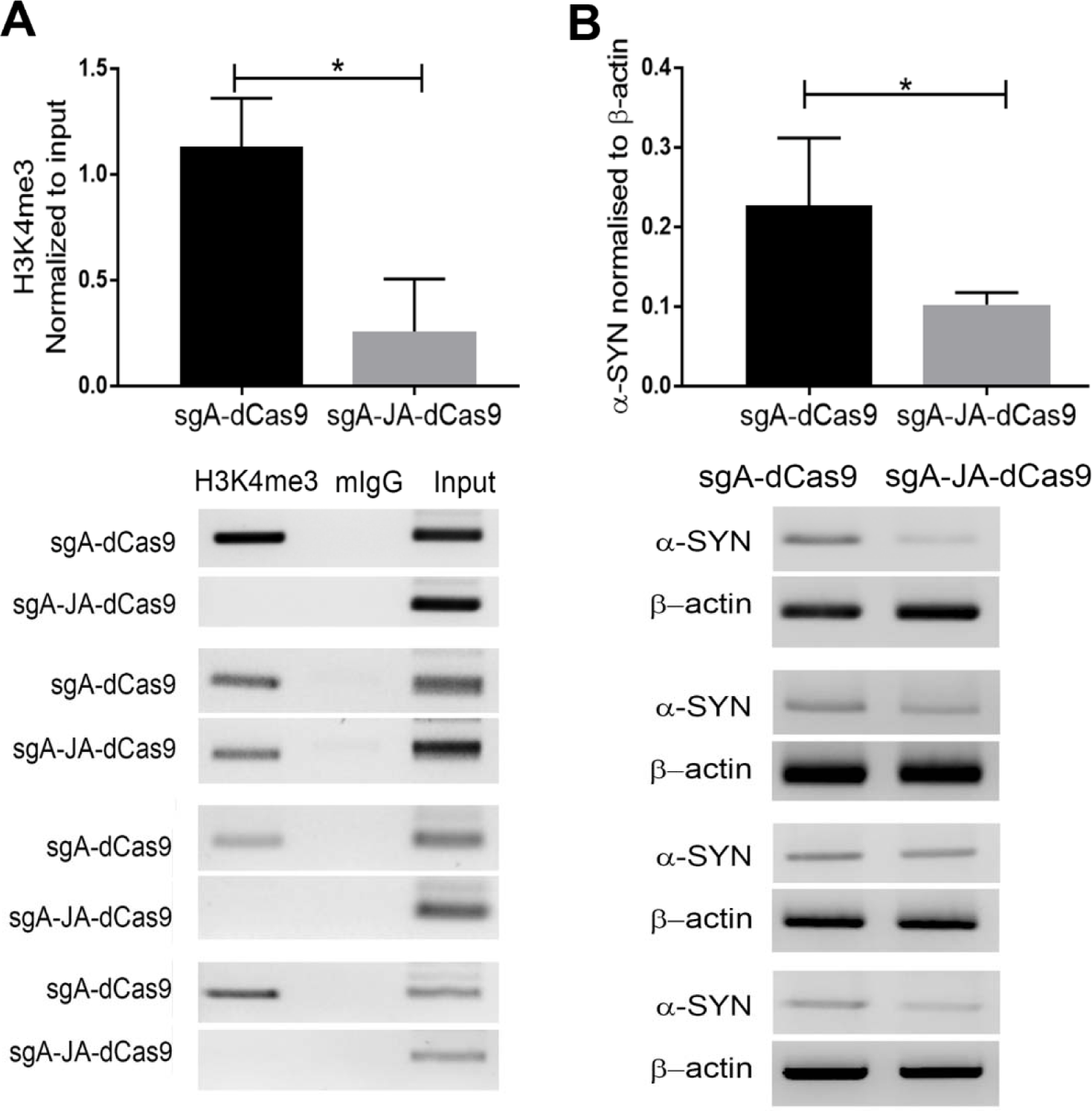
CRISPR/dCas9 SunTag-JARIDIA decreases α-synuclein expression by reducing H3K4me3 at the *SNCA* promoter. The relative enrichment of H3K4me3 (**A**) and corresponding α-synuclein levels (**B**) were evaluated in SH-SY5Y cells expressing dCas9-5xGCN4 with or without scFV-sfGFP-JARID1A. **A** ChIP data demonstrated a significant decrease of H3K4me3 at the *SNCA* promoter. The relative enrichment by H3K4me3 was normalized to the respective inputs. Four independent repeats were performed. **B** The levels of α-synuclein in SH-SY5Y cells were evaluated using RT-PCR. The level of α-synuclein was normalized to β-actin. A significant reduction of α-synuclein in cells expressing scFV-sfGFP-JARID1A was observed. Four independent repeats were performed. *\*P* < 0.05. Data were analyzed using non-parametric t-test followed by Mann-Whitney post-hoc corrections. Two-tailed p-values were calculated for both. Data information: All data are presented as mean ± SEM.

The sgA is located on non-coding exon 1B, which is approximately 1 kb upstream of the TSS (Fig 4C). SH-SY5Y cells expressing dCas9-5xGCN4/sgA/scFV-sfGFP-JARID1 A (dCas9-sgA-JA) exhibited significantly reduced H3K4me3 (almost to null) at the promoter of the gene (p<0.05) compared to control cells expressing dCas9-5xGCN4/sgA (dCas9-sgA) (Fig 5A). The significant reduction in H3K4me3 resulted in a notable decrease in expression of α-synuclein (Fig 5B). To confirm that reduction in α-synuclein was not caused by any of the individual components of the tripartite SunTag system, we separately overexpressed the three components in SH-SY5Y cells and did not observe any significant changes in α-synuclein levels (Fig EV8A). Next, we investigated global H3K4me3 levels in cell lines expressing dCas9-sgA-JARID1A compared to wild type SH-SY5Y cells. We did not observe any significant difference in global H3K4me3 levels between these cells (Fig EV8B).

### Efficient reduction in α-synuclein by CRISPR/dCas9 SunTag-JARIDIA in PD-patient derived iPSCs

We previously reported that dopaminergic neurons differentiated from sporadic PD patient-derived iPSC lines (sPD) expressed significantly higher levels of α-synuclein than did neurons derived from control iPSCs (Je et al, 2018). In addition, the level of tyrosine hydroxylase (TH) expression did not differ between PD and control iPSC lines when differentiated completely (Je et al, 2018). Therefore, we next investigated whether JARID1A could reduce α-synuclein levels in these sPD-iPSC lines. As expected, differentiated sPD-iPSC cells exhibited high enrichment of H3K4me3 at the *SNCA* promoter (Fig 6A). To determine if locus-specific recruitment of JARID1A could reduce H3K4me3 at the *SNCA* promoter in PD-iPSCs, we selected a sPD1-1 line and transiently transfected sgA with scFV-sfGFP-JARID1A and dCas9-5xGCN4 on day 25 of differentiation. We confirmed that sPD-iPSC lines were adequately differentiated by day 25-30; they exhibited sufficient matured dopaminergic neurons expressing TH (Fig EV9).

**Figure 6.**
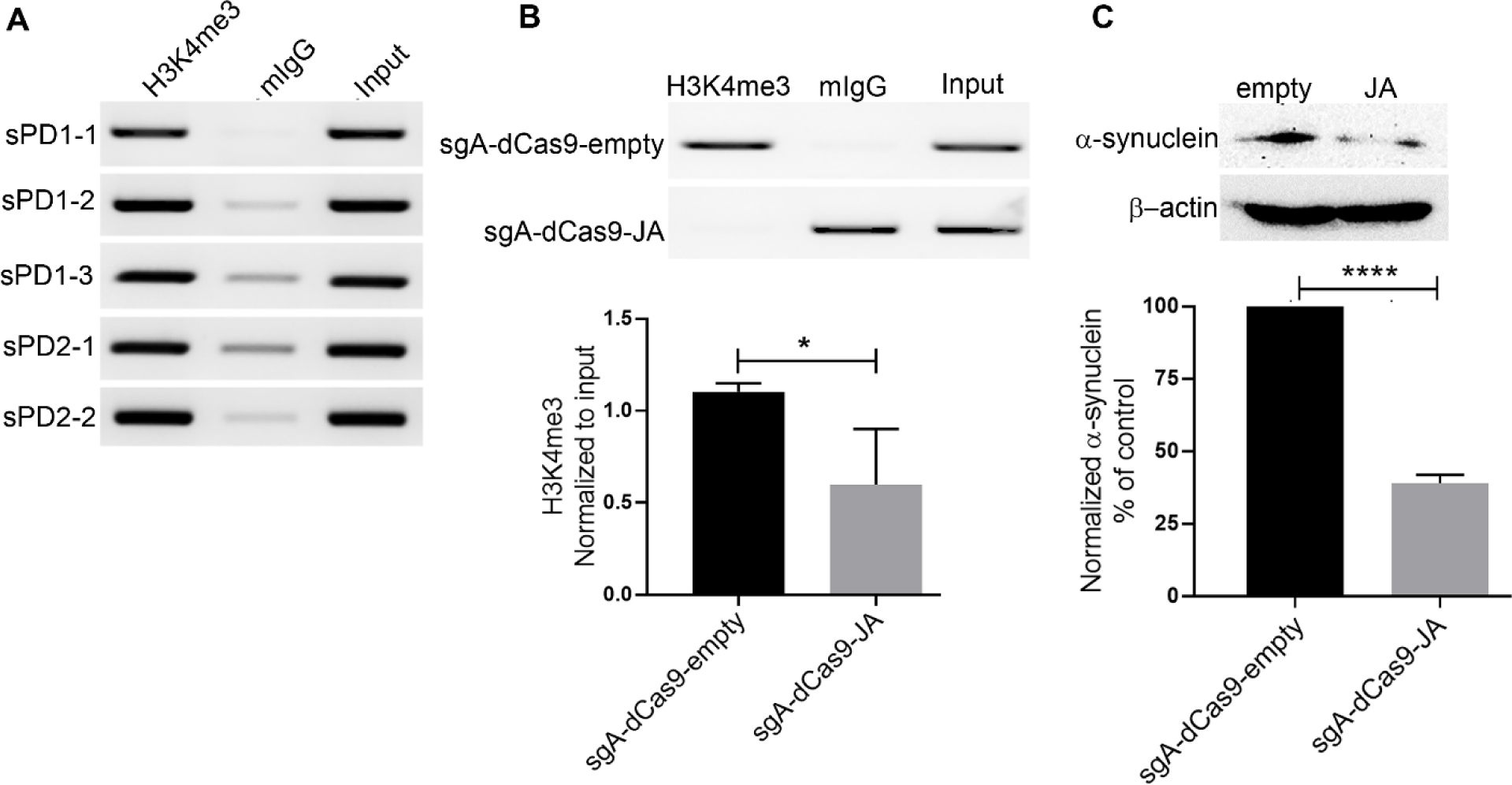
CRISPR/dCas9 SunTag-JARID1A ameliorates elevated α-synuclein levels in differentiated PD-iPSCs by reducing H3K4me3 from the gene promoter. **A** Relative enrichment of H3K4me3 at the *SNCA* promoter/intron 1 region was evaluated from five iPSC lines derived from two sporadic PD (sPD) cases by ChIP. The same genomic region as shown in Fig EV1 and Fig EV4 was evaluated for enrichment of H3K4me3. Mouse IgG (mIgG) was used as control for the target antibody. The relative intensity of the target band was normalized by respective input. **B** ChIP on H3K4me3 at the SNCA promoter in sPD1-1 line after locus-specific epigenomic modulation. Differentiated cells were transiently transfected with either sg A-dCas9 5xGCN4-scFV-JARID1A or sg A-dCas9 5xGCN4-scFV-empty backbone vectors. A significant reduction in H3K4me3 at the *SNCA* promoter was observed in cells transfected with JARID1A. A representative ChIP image is shown at the top. **C** Western blot analysis of α-synuclein levels in the cells under the same conditions as analyzed in (B). A significant decrease (56-66%) in α-synuclein levels was observed in cells transfected with JARID1A. The normalized and relative expression of α-synuclein in JARID1A transfected cells are shown as a percentage of control (transfected by empty backbone vector). A representative western blot image is shown at the top. Three independent repeats were performed for both ChIP and western blot experiments. *\*P* < 0.05. Data were analyzed using non-parametric t-test followed by Mann-Whitney post-hoc corrections. One-tailed p-values were calculated for ChIP analysis while two-tailed p-value was calculated for western blot experiments. Data information: All data are presented as mean ± SEM.

Successful decrease in H3K4me3 at the *SNCA* promoter was achieved after transfection with dCas9-5xGCN4/sgA/scFV-sfGFP-JARID1A compared to control (dCas9-5xGCN4/sgA/scFV-empty) (Fig 6B). As expected, we observed very significant reduction (average decrease of 56-66%; p<0.00001) in α-synuclein protein levels in differentiated sPD-iPSC lines administered with the CRISPR/dCas9 SunTag-JARID1A system (Fig 6C). These results indicate that locus-specific modulation of H3K4me3 at the *SNCA* promoter effectively ameliorated α-synuclein in PD patient-derived iPSC lines.

## Discussion

Properly orchestrated regulation by epigenetic factors ensures successful gene expression; any disparities in epigenetic regulation can lead to faulty transcription. *SNCA,* the gene encoding α-synuclein, has a complex epigenetic environment and it is reasonable to hypothesize that deregulation could lead to altered gene expression (Guhathakurta et al, 2017a). In this study, we performed an in-depth analysis of the epigenetic regulation of *SNCA* in PD. We demonstrated that H3K4me3, a histone mark associated with transcription initiation, could serve as a pivotal epigenetic marker for the deregulated expression of *SNCA* in PD. Our data using postmortem brain samples from PD and matched controls clearly indicate that H3K4me3 is significantly enriched at the *SNCA* promoter/regulatory region in PD patients. Successful locus-specific reduction of this histone mark from the *SNCA* promoter reduced levels of α-synuclein significantly in human neuroblastoma, SH-SY5Y, and PD patient-derived iPSC cell lines. This finding could open new research avenues for development of therapeutic strategies for PD and other neurodegenerative diseases by targeting the epigenetic environment of implicated genes.

The promoter and associated regulatory region of human *SNCA* contain several histone PTMs that can regulate α-syncul ein’s transcriptional state. Three histone PTMs that were enriched at the *SNCA* promoter were H3K4me3, H3K27ac, and H3K27me3. While both H3K27ac and H3K4me3 were occupied at the *SNCA* promoter, only H3K4me3 was found to be enriched significantly in the patient group compared to controls. We did observe H3K27me3 was preferentially enriched in a subset of patients; however, this enrichment was not as significant as H3K4me3. This selective and most significant enrichment of H3K4me3 at the *SNCA* promoter might account for higher expression of α-synuclein in the SN of PD patients. As mentioned in the methods, PCR cycles were kept to 35 for these ChIP-PCR studies to show the maximum difference of SNCA promoter-specific enrichment by H3K4me3 between control and PD subjects. control subjects did show some degree of H3K4me3 enrichment when the PCR cycles were increased to 40 cycles where saturation was observed in input.

Brain tissue has complex architecture comprising different cell types specific to regions of the brain (Guintivano et al, 2013). A significant difference in enrichment of H3K4me3 at the *SNCA* promoter/intron 1 region was initially observed using entire tissue sections from the SN. Because the presence of heterogeneous cellular populations in the region can confound results, epigenetic environment can vary by cell type (Guintivano et al, 2013), and interpretation of data from the ensemble average of SN tissue for a disease characterized by selective dopaminergic neuronal loss is potentially risky, we further investigated whether the epigenetic changes observed in the entire tissue composition were also present in pure neuronal populations from the SN region. We examined H3K4me3 at the *SNCA* promoter/intron 1 from NeuN+ neurons isolated from SN tissue of a subset of study subjects. NeuN-FANS yields a homogenous population of nuclei from neurons. The equal number of neuronal nuclei isolated from both groups also showed a significant H3K4me3 enrichment at the *SNCA* promoter in the patient samples, suggesting that SN neurons in PD patients suffer from a significant imbalance in epigenetic regulation of α-synuclein. A technical limitation, however, made it difficult to investigate epigenetic changes selectively in dopaminergic neurons. We also observed significantly higher amounts of α-synuclein protein in the same SN region in PD compared to controls. However, not all the patients exhibited equally high levels of α-synuclein, nor did the controls all exhibit low levels of α-synuclein. Therefore, to determine if high levels of α-synuclein were correlated with H3K4me3 enrichment, we segregated the entire cohort into high and low α-synuclein expressing groups. Overall, high levels of α-synuclein were significantly correlated with higher enrichment of H3K4me3 at the *SNCA* promoter. However, this correlation did not follow complete linearity, suggesting other secondary epigenetic factors could be involved in this complex gene regulation.

Subsequently, we explored two other major histone marks in the same region of the gene, H3K27ac and H3K27me3. The transcription favoring histone mark, H3K27ac, did not show any bias toward PD patients, while the transcription repression mark, H3K27me3, was increased in PD patients compared to controls. Although significant, enrichment of H3K27me3 in patients was obviously lower than relative enrichment of H3K4me3 in the same patients. The relatively higher levels of H3K27me3 and low levels of H3K27ac at the *SNCA* promoter in PD patients might counterbalance the effect of H3K4me3, leading to moderately significant levels of α-synuclein in PD patients. It is well known that H3K27ac and H3K27me3 compete for the same lysine residue on an H3 peptide tail. Therefore, coexistence of these two marks in a single nucleosome is relatively unlikely. We also have observed that levels of H3K27ac are relatively higher in SN tissues (with no difference between PD and controls) as compared to occupancy by H3K27me3 at the promoter of *SNCA.* Epigenetic analysis of a single neuron could reveal whether an individual neuron in the SN has differential epigenetic markers, such as H3K4me3/H3K27ac or H3K4me3/H3K27me3. This specific combination of epigenetic architecture could be a cell-type specific event. Previously, we and other groups independently showed that *SNCA* has the intragenic enhancer region represented by the enrichment of H3K27ac in intron 4 (Guhathakurta et al, 2017a; Soldner et al, 2016). However, we failed to observe significant difference between the groups in enrichment of this enhancer-associated histone mark.

Next, we aimed to determine whether locus-specific removal of H3K4me3 could reduce pathological protein levels. To achieve selective reduction in H3K4me3 enrichment from the *SNCA* promoter, we made a genomic locus-specific molecular tool employing a unique H3K4me3 erasing enzyme, JARID1A. Locus-specific epigenomic editing using the dCas9-CRISPR technology has been shown to be efficacious and powerful in changing transcriptional states of target genes. Several different models have been developed, and the majority of them employ general transcriptional activators, such as VP64, p65, or Rta (or a combination of all three), and synergistic activation mediator (SAM; MS2-p65-HSF1)-based systems to activate gene expression by modulating local chromatin structure (CRISPR-activator) (Chavez et al, 2015; Chavez et al, 2016; Gilbert et al, 2013; Konermann et al, 2015; Tanenbaum et al, 2014). Interestingly, it has also been shown that fusing a catalytic core domain of histone acetyl transferase (p300) to dCas9 and targeting it to the enhancer of the target genes could activate transcription of multiple genes (Hilton et al, 2015). Simultaneously, transcriptional inhibition systems (CRISPR-inhibitor) were developed by recruiting various transcriptional inhibitory proteins such as KRAB (Kruppel associated box domain of Kox1) or four concentrated copies of mSin3 interaction domain (SID4X) within a few hundred bases downstream of the TSS of the target genes (Gilbert et al, 2013; Konermann et al, 2013). These systematic genetic interrogation systems have proven extremely beneficial in whole genome screening platforms for identifying disease-phenotype related genes and genes responsible for survival against toxins and chemotherapeutic drugs (Gilbert et al, 2014; Shalem et al, 2015). However, only a handful of studies have investigated the beneficial effects of altering the epigenetic environment of disease-implicated genes in disease models using epigenetic writer enzymes (Choudhury et al, 2016; Liu et al, 2018).

To edit the local epigenetic structure of the *SNCA* promoter most efficiently, we used the CRISPR/dCas9 SunTag system where dCas9 tagged with tandem repeats of an epitope allowed us to bring multiple effector molecules to the target site to enhance its effect (Tanenbaum et al, 2014). First, we redesigned the original SunTag system that was developed to recruit 10 to 24 copies of VP64 to the target site using a single guide RNA, achieving significant transcriptional activation of the target genes (Tanenbaum et al, 2014). JARID1A enzyme was selected to remove H3K4me3. Previously, Hilton *et al*. showed that using the catalytic core of JARID1A as an epigenetic eraser works more efficiently than using the whole enzyme (Hilton et al, 2015). Since it was previously shown that first 797 amino acids of JARID1A has the most capacity to remove H3K4me3, we replaced VP64 in the original SunTag system with the catalytic domain of JARID1A. Considering the relatively large size of the JARID1A catalytic domain, to avoid steric hindrance and proper recruitment of this bulky enzyme, we added 22-amino acid spacers in between five GCN4 tails at the c-terminus of dCas9 (Morita et al, 2016). Our modified system was able to reduce significantly enrichment of H3K4me3 at the *SNCA* promoter when stably overexpressed in SH-SY5Y cells. For the target region on the *SNCA* promoter/intron, we were not restricted to recruit effectors to -200 to -400 bp of the TSS since we were not modulating gene expression directly using transcriptional activators or inhibitors. Instead, we designed guide RNAs based on the distribution of H3K4me3 peaks upstream of the TSS in *SNCA* (from -132 bp to -1537 bp). We observed that in SH SY5Y cells, targeting the area around 1 kb upstream of the TSS most significantly reduced H3K4me3 from the promoter. As shown in Fig 5B, the reduction of H3K4me3 from those sites in *SNCA* significantly reduced α-synuclein levels, indicating that H3K4me3 is one of the principal epigenetic regulators of the gene. CRISPR/dCas9 SunTag-JARID1A is a tripartite system requiring all three components, sgRNA, dCas9-5xGCN4, and scFV-sfGFP-JARID1 A, to ensure removal of H3K4me3 mark from the target site.

Overexpression of any of the individual components did not result in any change in α-synuclein levels, demonstrating that the effect on α-synuclein was due to focal recruitment of SunTag-JARID1A at the *SNCA* promoter. Minimal off-target reduction of H3K4me3 was observed; we did not find any changes in global H3K4me3 levels in the genome when we compared relative levels of global H3K4me3 between wild-type cells with sgA-dCas9-JARID1A line. Previously, Morita *et al*. reported that the dCas9-based SunTag system with a TET1catalytic domain as an effector molecule had exceptionally low off-target effects using whole genome bisulphite sequencing (Morita et al, 2016). Moreover, when Kantor *et al*. also targeted dCas9-DNMT3A at the *SNCA* intron 1 to increase methylation, they did not find any significant change in global DNA methylation (Kantor et al, 2018).

After ensuring that the CRISPR/dCas9 SunTag-JARID1A system could efficiently reduce H3K4me3 from the *SNCA* promoter without perturbing epigenetic architecture elsewhere in the genome, we studied its effect on idiopathic PD-derived iPSC lines. As we reported previously, differentiated dopaminergic neurons from these lines exhibited high levels of α-synuclein compared to control lines, which correlated well with the H3K4me3 levels at the gene promoter. We next investigated whether the locus-specific approach was similarly effective in reducing α-synuclein in PD-iPSC lines after differentiation. Since iPSC lines are known to suppress any transgene promoter activity, it was extremely difficult to make stable cell lines expressing all components of the CRISPR/dCas9 SunTag-JARID1A system (Norrman et al, 2010). Since the entire system is bigger than the lentiviral genome, we could not use an “all-in-one” lentiviral system. Instead, we adopted an efficient transient transfection method using magnetofection. This magnet-based co-transfection ensures high transfection efficiency in primary dopaminergic neurons (Underhill et al, 2014). We also observed a good transfection rate in differentiated dopaminergic neurons derived from the PD iPSC lines. As expected, CRISPR/dCas9 SunTag-JARID1A again significantly reduced H3K4me3 at the *SNCA* promoter with a concomitant decrease in α-synuclein levels of around 55-66 percent. Previously, in an elegantly designed study, Soldner *et al*. showed that editing a PD-associated point mutation at the intronic enhancer region of *SNCA* could affect transcription factor binding and transcription efficiency (Soldner et al, 2016). However, this mutation is rare in the population and it was shown that it could affect expression of α-synuclein up to 1.18 times in cell culture models. Another recent study by Kantor *et al*. demonstrated significant reduction of α-synuclein by recruiting dCas9-DNMT3A (a single copy of DNMT3A directly fused at the end of dCas9) to *SNCA* intron 1 in an *SNCA*-Tri hiPSC line derived from dopaminergic progenitor cells (Kantor et al, 2018). Although the results were significant, it should be noted that several groups did not find any difference in DNA methylation between PD patients and matched controls (Guhathakurta et al, 2017b). Therefore, increasing methylation at *SNCA* intron 1 might not be physiologically relevant. Moreover, as reported previously (Gilbert et al, 2013), recruiting dCas9-DNMT3A downstream to the TSS itself might directly hinder transcription by disrupting progress of polymerase along the gene. In our study, we avoided this issue by targeting the region upstream of the principal TSS of the gene where H3K4me3 was found to be enriched.

In sum, our study describes a novel and efficient method of modulating deregulated expression of α-synuclein in PD. Despite its extreme importance, not many studies have investigated epigenetic control of *SNCA* in neuronal cells or dopaminergic neurons. With advanced molecular techniques like CRISPR/Cas9 and advanced knowledge of tissue-specific epigenetic environment of all genes, it is now possible to edit the local genetic and epigenetic microenvironment of genes implicated in diseases. Here, we reported significant epigenetic deregulation of the *SNCA* gene in PD and a novel strategy that efficiently ameliorated expression of α-synuclein by editing pathological epigenetic marks in differentiated iPSCs derived from idiopathic PD patients. The study indicates a potential therapeutic application in PD by epigenetic modulation of H3K4me3 at *SNCA,* which might ameliorate α-synuclein-mediated degenerative changes in the disease. However, future studies are necessary to investigate whether reduction of α-synuclein by epigenetic editing could rescue degeneration in PD using animal models or iPSC-based models of PD.

## Materials and methods

### Brain tissues

All human postmortem brain samples were obtained from NIH Neurobiobank as mentioned in our previous publication (Guhathakurta et al, 2017b). In this study, we have increased our patient cohort from 8 samples (as previously reported (Guhathakurta et al, 2017b)) to 18 samples. All samples were ethnicity-, age-, and sex-matched. Ages ranged from 70 to 89 years for the PD cohort and 54 to 89 years for controls. Postmortem interval (PMI) ranged from 10.1 to 35.42 hours in PD and 10.0 to 30.25 hours in controls. The PMIs and the age ranges did not vary significantly between groups. Details of postmortem brain samples are provided in Supplementary Table 1.

### Antibodies

The following antibodies were used for western blot analysis: α-synuclein (BD Transduction Laboratories, Clone 42/α-synuclein (RUO), 610787; dilution 1:500); β-actin (Sigma, A5316; dilution 1:20,000); Flag-tag (Sigma, F3165; dilution 1:5,000); Horseradish peroxidase (HRP) conjugated **s**econdary (goat anti-mouse IgG, Jackson Laboratory, 115-035-146; goat anti-rabbit IgG, Jackson Laboratory, 111-035-144; 1:5000 dilution). For FACS analysis, anti NeuN antibody was used (EMD Millipore, ABN78; 1:500). For chromatin immunoprecipitation experiments, the following primary antibodies were used: (Normal mouse IgG, Millipore, 12-371; H3K4me3, Abcam, ab8580; H3K27me3, Active motif, 39155; H3K27ac, Abcam, ab4729). Anti-Cas9 antibody (Takara, 632607; 1:1000) and anti-GFP antibody (Fisher Scientific, MS1315P0; 1:5000) were used for immunoprecipitation experiments. For immunocytochemistry, TUJ1 (Neuromics, MO15013; 1:500) and TH (Santa Cruz, SC-25269; 1:200) antibodies were used.

### Cell culture

SH-SY5Y cells were purchased from ATCC and maintained as per their guidelines. Briefly, cells were grown in DMEM/F12 medium (Thermo Scientific, SH30023FS) containing 10% FBS. The HEK293T cells were cultured in DMEM/high glucose medium (Thermo Scientific, SH30243FS) supplemented with 10% Fetal Bovine Serum (FBS) (Atlanta Biologicals, S10350H). The cells were maintained in a humidified atmosphere of 5% CO_2_ at 37°C. Cells were seeded at a density of 1.2 x 10^6^ cells/10 cm dish, 7.5 x 10^5^ cells/well of 6-well plates and 0.5 x 10^5^ cells/well of 24-well plates.

### iPSC culture and differentiation of PD-iPSC derived dopaminergic neurons

Five iPSC lines derived from two PD patients used in the study were provided by Dr. Hanseok Ko at the Johns Hopkins University School of Medicine. The iPSC lines were differentiated into dopaminergic neurons following a previously published protocol with some modifications (Kriks et al, 2011). PD-iPS cells were expanded on Matrigel in mTeSR1 media (Stemcell Technologies, 85850) with doxycycline (1 μg/ml, Sigma, D9891) (Chang et al, 2014). Cells were dissociated into a single-cell suspension using accutase and seeded at a density of 1.0 x 10^6^ cells per well on Matrigel coated 6-well plates in mTeSR1 supplemented with 10 mM ROCK inhibitor Y-27632 (Tocris, 1254).

For differentiation, cells were exposed to 100 nM LDN193189 (Reprocell, 04-0074) from 0∼10 days, 10-μM SB431542 (Reprocell, 04-0010) from days 0-6, 100 ng/ml recombinant human sonic hedgehog (SHH) (R&D systems, 464-SH-025CF), 2 μM purmorphamine (Reprocell, 04-0009), and 100 ng/ml recombinant human/mouse FGF-8b (Peprotech, 100-25) from days 1-6 and 3-μM CHIR99021(Reprocell, 04-0004) from days 3-12. Cells were grown for 11 days in knockout serum replacement medium (KSR; Gibco, 10828-028) containing advanced DMEM/F12 (Gibco, 12-634-010), 20% KSR, 2-mM L-glutamax (Gibco, 35050-061), and 10-mM β-mecaptoethanol (Gibco, 21985-023). The ratio of KSR medium to N2 medium started at 75:25% and was gradually changed to 25:75% starting from day 5 of differentiation to day 10. On day 11 of differentiation, media was changed to B27 media containing Neurobasal media (Gibco, 21103-049), B27 serum supplement (Gibco, 17504-044), and L-glutamax supplemented with CHIR (3 μM), 20 ng/ml BDNF(PeproTech, 450-02), 20 ng/ml GDNF(PeproTech, 450-10-50ug), 0.2 mM ascorbic acid (Fisher Scientific, A61-100), 1 ng/ml TGFb3(R&D Systems, 243B3002/CF), and 0.1 mM dibutyryl cAMP (Sigma, D0627-250MG). From day 13 to 20, the same media composition was used except CHIR was added.

On day 20, cells were dissociated using accutase and replated under high cell density conditions (1x10^5^ - 3x10^5^ cells per cm^2^) on dishes pre-coated with 15 μg/ml polyornithine, 5 μg/ml laminin, and 2 μg/ml fibronectin in differentiation medium for 10∼15 days using day 20 media changed alternating days.

### Plasmid constructs

All template plasmids were sourced from Addgene. Specifically, the pCAG-dCas9-5xPlat2AflD plasmid (Addgene #82560) was a gift from Izuho Hatada (Morita et al, 2016). The lentiGuide-puro (Addgene #52963) and pHRdSV40-scFv-GCN4-sfGFP-VP64-GB1-NLS (Addgene #60910) plasmids were gifts from Feng Zhang (Sanjana et al, 2014) and Ron Vale (Tanenbaum et al, 2014), respectively. The catalytically active domains of JARID1A enzyme (1-797 amino acids) was amplified from pcDNA3/HA-FLAG-RBP2 plasmid (Addgene #14800), a gift from William Kaelin (Klose et al, 2007). The amplified fragment containing linker sequences along with *RsrII* sites at both ends was then sub cloned into pHRdSV40-scFv-GCN4-sfGFP-VP64-GB1-NLS plasmid after removing the VP64 fragment by restriction digestion using *RsrII*. To generate the empty backbone vector, the pHRdSV40-scFv-GCN4-sfGFP-VP64-GB1-NLS plasmid was digested with *RsrII* enzyme and ligated back to compatible ends of the restriction enzyme. The backbone vector produced this way retained the entire promoter—scFV, sfGFP, NLS and GB1 components (only VP64 was absent). To generate lentivirus from these constructs, all of them were transferred to pLvx-DsRed vector backbone (Clontech). The entire ScFv-sfGFP-JARID1A-GB1-NLS sequence was sub-cloned into pLvx-DsRed vector using *XmaI/SexAI* restriction sites. Using this strategy, we removed the DsRed, PGK promoter, and puromycin from the pLvx vector and replaced in frame with scFV-JARID1A. Importantly, the original Addgene plasmid (#60910) was also a lentiviral vector, but we sub-cloned the insert into pLvx-DsRed vector because the Addgene vector lacked a WPRE sequence, which resulted in low transduction efficiency. Similarly, dCas9-5X GCN4 was also sub-cloned into pLvx-DsRed vector at *XmaI/NotI* restriction sites.

All guide RNAs targeting the *SNCA* promoter were cloned into LentiGuide-puro vectors following a previously described protocol (Sanjana et al, 2014). The guide RNAs were designed using the Broad Institute genetic perturbation platform (https://portals.broadinstitute.org/gpp/public/analysis-tools/sgrna-design) and were re-verified using the CHOPCHOP program (https://chopchop.cbu.uib.no/). To increase specificity and reduce off-target effects, at least 3 mismatches were allowed at the 3’end before the PAM sequence.

### Transfection

All transient transfection was carried out using magnetofection protocol by Oz Biosciences using their Neuromag reagent. Briefly, differentiated iPSC cells were grown in a 12-well plate and the media was changed on the day of transfection with 1 ml of antibiotic free media. The cells were transfected first on day 25 of differentiation. Around 3 μg of total plasmid DNA was transfected per well following equal molar mass for all three vectors (0.16 pmols each). The DNA was then diluted in 150 μl of Opti-MEM media and spun down. The diluted DNA was mixed gently with 9 μl Neuromag reagent in a different tube and allowed to incubate for 30 minutes before adding to the cells. After adding the transfection mix to the cells, the plate was put on a magnetic bar overnight in the 37°C incubator and the next day the magnet was removed underneath the plate. This process ensured a very high transfection rate without any visible cell mortality. Transgene expression was visualized by expression of sfGFP associated with the SunTag system. To increase transfection efficiency, cells were transfected again the same way after 3 days of the first transfection and harvested 6 days after first transfection.

### Establishing stable SH-SY5Y cell lines expressing the SunTag system

To obtain a pure enrichment by sgRNA and dCas9, cells were treated serially with lentiviral particles containing dCas9 5xGCN4 and sgRNA. The sgRNA vector was resistant to puromycin and dCas9 was resistant to blasticidin. To generate high titer lentiviral particles from the three components of the SunTag system, the individual transfer plasmids were mixed with lentiviral packaging plasmids, psPAX2 and pMD2.G, at a 1:1:1 molar ratio using X-fect transfection reagent in HEK293T cells. Around 48 hours post transfection, the medium containing lentiviral particles was filtered through 0.45 uM PES filter and mixed with lenti X concentrator at a ratio of 1:4. The solution was then incubated at 4°C for a minimum of 4 hours and centrifuged for 45 minutes at 1,500 X g to pellet down the viral particles. The pellet was then resuspended in an appropriate amount of sterile phosphate buffered saline (PBS) to bring the final concentration to 100X. To generate SH-SY5Y cells containing dCas9-5xGCN4, 10 μL concentrated viral particles mixed with 4 μg/ml polybrene (Sigma) was added to the SH-SY5Y cells grown on a 24-well plate. After 48 hours of treatment with lentiviral particles, the cells were positively selected under antibiotic blasticidin (5 μg/ml) (Acros Organics, 227420100) for 4 days. The cells that grew under antibiotic selection were expanded and received a second lentiviral transduction for sgRNA. Cells were transduced with 10 μl sgRNA containing lentiviral particles and were allowed to grow for another 2 days before selection with puromycin (2 μg/ml) for 48 hours. The puromycin resistant cells were expanded for 1 week. Cells were then treated with JARID1A containing lentiviral particles for 72 hours. Finally, to obtain a pure population of cells containing all three transgene expressions, cells were FAC sorted for GFP (conferred by JARID1A/backbone plasmid). Sorted cells were then grown for another 2 days before harvesting for biochemical analysis.

### Fluorescence activated nuclei sorting (FANS)

FANS from postmortem brain samples were conducted following published protocols with some modifications (Jiang et al, 2008). Briefly, the frozen postmortem brain tissue (100 mg) around the SN region was dissected on a cryotome. The tissue was then transferred to the glass container of Dounce homogenizer, which already contained 5 ml Nuclear Extraction Buffer (NEB; 0.32 M Sucrose, 5 mM CaCl2, 3 mM Mg(Ac)2, 0.1 mM EDTA, 10 mM Tris-HCl (pH8), 1 X Protease inhibitor cocktail, 0.1% Triton X-100) with 1% formaldehyde. Once the tissue was thawed in NEB, it was homogenized 21 times with loose pestle and 9 times with tight pestle for about 1 minute (Hu et al, 2017). After homogenization, 5 ml of sample solution was transferred to a 15-ml tube at RT for 10 minutes for optimum crosslinking. Excess formaldehyde was quenched by adding 500 μL 125 mM glycine for 5 minutes. The homogenized sample was then pelleted by centrifuging it at 2,000 X g for 5 minutes. The supernatant was discarded and the pellet was resuspended in 5 ml fresh NEB supplemented with protease inhibitor (PI). The resuspended nuclei were then transferred to 38.5-ml ultracentrifuge tube on ice (Beckman Coulter, 344058). To the nuclei suspension, 25 ml of sucrose cushion buffer (1.8 M Sucrose, 3 mM Mg(Ac)2, 10 mM Tris-HCl; pH 8.0) was added to the bottom of the tube. Any volume make-up to the top of the tube were done using NEB. The samples were then centrifuged at 107,163.6 X g for 2.5 hours at 4°C in an ultracentrifuge (XPN 100, Beckman Coulter). After centrifugation, the supernatant was removed from the tubes on ice and 1 ml pre-chilled PBS was added to the nuclei pellet. The tubes were incubated for 20 minutes on ice without any disturbance and the pellet was resuspended optimally. The sample was then transferred to a tube and mixed with nuclei resuspension buffer (NRB: 250 mM sucrose, 25 mM KCl, 5 mM MgCl2, 20 mM Tri-Cl, 1.0%. BSA, pH 7.4; Supplemented with PI) at 1:2 ratio. The nuclei were then recovered after 5 minutes centrifuge at 2,000 X g and again resuspended in 1 ml NRB on ice. The integrity of the nuclei and number was checked under the microscope from a small aliquot of the resuspended nuclei. From that 1 ml of nuclei suspension, 100 jul suspension was aliquoted in a different tube to make appropriate negative controls such as unlabeled or unstained controls. The volumes were brought to 1 ml with NRB. To the remaining 900 μl nuclei, 1 μl rabbit anti-NeuN antibody was added and the volume was brought to 1 ml with NRB. All the tubes were then incubated overnight at 4°C on a rotating platform. The next day, nuclei were recovered by spinning down at 2,000 X g for 5 minutes and washed twice for 10 minutes with NRB before incubating with secondary antibody. To the washed sample, 1 μl goat anti-Rb IgG alexa Fluor-488 secondary antibody was added and incubated for 1 hour on a rotating platform at RT in the dark. Next, samples were again washed twice with NRB for 10 minutes each before FAC sorting. Finally, the samples for FANS were resuspended in PBS+PI and strained through 40-μM nylon filter right before sorting. FANS gatings were made based on a previously published protocol (Jiang et al, 2008). Samples prepared for chromatin immunoprecipitation (ChIP) reactions were directly collected in 350 μL SDS lysis buffer (composition is mentioned under ChIP protocol). A Biorad 2 laser S3e sorter was used to sort the samples.

### Chromatin Immunoprecipitation (ChIP)

ChIP was performed following the protocol for EZ ChIP™ Chromatin Immunoprecipitation kit (Millipore, 17-371) with the following modifications. The treated or untreated adherent cells (SH-SY5Y and differentiated iPSCs) were fixed by adding required volume of 37% formaldehyde (final concentration 1%) to the medium and incubated for 5 minutes at RT. The excess formaldehyde was quenched using 125-mM glycine for 10 minutes. Cells were collected in PBS (pH 7.4) containing PI cocktails following two washes in PBS. For human postmortem SN tissues, 22 mg of freshly frozen tissue from each sample was isolated by punch biopsy on dry ice and homogenized by Teflon coated homogenizer pre-cooled in liquid nitrogen. The sample was then cross-linked in 1 ml 1% formaldehyde for 20 minutes at RT. The supernatant was discarded following a brief spin and 125-mM glycine was added for 5 minutes to each sample.

The samples were then washed twice in 1 ml PBS supplemented with PI. After this step, the tissues were processed as described below. Collected cells/tissues were centrifuged at 400 X g for 3 minutes and resuspended in 350 μl lysis buffer (1% SDS; 10-mM EDTA, pH 8.0; 50-mM Tris-Cl, pH 8.0) for 10 minutes on ice. Cell suspension was then sonicated in a sonicator (Fisher Scientific, Model No. FB 50) equipped with a probe for micro-centrifuge tubes (Model No. CL-18, Fisher Scientific) for 5 pulses at 20 Hz for 20 seconds with 30 second intervals in between each pulse. The sonicated cells were centrifuged at 10,000 X g for 10 minutes at 4°C. Remaining supernatant was collected and diluted 10X in ChIP dilution buffer (0.01% SDS; 1.1% Triton X-100; 1.2-mM EDTA, pH 8.0; 16.7-mM Tris-Cl, pH 8.1; 167-mM NaCl) containing PI. Diluted chromatin was pre-cleared with Salmon Sperm DNA/Protein A agarose slurry (Millipore, 16-157) at 50 μl slurry/ml of diluted chromatin in an end-to-end rotator overnight at 4**°**C. The next day in the morning, 30 μL pre-cleared chromatin was kept as input and 900 μl chromatin was incubated with each target antibody (1 μg) at 4°C in a rotor. In addition, 20 μL Salmon Sperm DNA/Protein A agarose slurry was added to each tube after 8 hours of incubation with the antibody and left for a combined incubation at 4°C overnight. The next day, the immune complex was retrieved by centrifuging at 4,000 X g for 1 minute and sequentially washed 1 time in low salt wash buffer (0.1% SDS; 1.0% Triton X-100; 2-mM EDTA, pH 8.0; 20-mM Tris-Cl, pH 8.1; 150-mM NaCl), 1 time in high salt buffer (0.1% SDS; 1.0% Triton X-100; 2-mM EDTA, pH 8.0; 20-mM Tris-Cl, pH 8.0; 500-mM NaCl), 1 time in Lithium Chloride (LiCl) wash buffer (250-mM LiCl; 1.0% IGEPAL; 1-mM EDTA, pH 8.0; 10-mM Tris-Cl, pH 8.1; 1.0% Deoxycholic acid), and finally washed twice in Tris-EDTA buffer, pH 8.0 (10-mM Tris-Cl; 1-mM EDTA). The antibody conjugated with fragmented DNA was then eluted by incubating it twice in 0.1-mM NaHCO3; 1.0% SDS buffer for 15 minutes each. Reverse crosslinking was initiated by adding 200-mM NaCl to the buffer and incubated overnight at 65°C. The next day it was incubated with 10 μg RNase A (Thermo Fisher Scientific) for 30 minutes at 37°C to remove RNA contamination. Completion of reverse crosslinking was achieved by adding 10-mM EDTA, pH 8.0; 40-mM Tris-Cl, pH 8.0; 50 μg Proteinase K and incubated at 45°C for 2 hours. The fragmented DNA was isolated by phenol: Chloroform: Isoamyl alcohol, DNA pellet was visualized by adding 5-10 μg glycogen and finally dissolved in 20 μl of DNase/RNase free water. ChIP DNA was then amplified for a segment (188 bp) of *SNCA* intron 1 using primer pair SNCA F3/R3. The sequence is presented in Table S2. The intensity of the band amplified from mouse IgG control was subtracted from each amplicon from respective target of IP’d DNA and normalized by 1% input from each sample.

For ChIP performed from FANS samples, equal number of GFP+ nuclei were collected in 350 μl SDS lysis buffer supplemented with PI. They were immediately sonicated following previously described parameters. Next, the volume was brought to 2 ml with ChIP dilution buffer, 100 μl protein A/G agarose beads, and PI. The remainder of the process was done as described above.

### Immunocytochemistry

To stain the differentiated dopaminergic neurons from PD-derived iPSC lines, a standard immunocytochemical method was followed (Cristovao et al, 2012). Briefly, cells were allowed to differentiate on coated coverslips in a 24-well plate. On day 30 of differentiation, cells were fixed and permeabilized with 4% paraformaldehyde in PBS-T (0.01% tween-20) and incubated overnight at 4°C with primary antibodies (TUJ1, TH) at indicated concentrations. The next day, cells were washed with wash buffer (PBS-T) several times and incubated with appropriate fluorophore labelled secondary antibodies (Alexa fluor 547) at indicated concentrations at RT for one hour in the dark. Cells were washed to eliminate non-specific binding and incubated with DNA labeling dye Hoechest132 (20 uM) before mounting on glass slides. Slides were visualized on a Nikon Ti eclipse inverted fluorescence microscope.

### Western blot

Cells grown on 10-cm dish or 6-well plates were harvested in cold PBS and pelleted down by centrifugation at 600 X g for 5 minutes at 4°C. Pelleted cells were subjected to lysis in an appropriate volume of Radio Immuno Precipitation Assay buffer (RIPA; 1% NP-40; 0.5% Sodium Deoxycholate; 0.1% SDS) supplemented with 1X protease and phosphatase inhibitor (Thermo Scientific, 1860932) for 15 minutes on ice, and supernatant containing proteins were collected following centrifugation at 12,000 X g for 15 minutes at 4°C. Approximately 40 μg protein sample was prepared in sample buffer (50-mM Tris-HCl, pH 6.8; 2% SDS; 10% glycerol; 1% β-mercaptoethanol; 12.5-mM EDTA; 0.02% Bromophenol blue) and boiled for 5 minutes followed by snap chilling on ice. The samples were loaded onto 10% SDS gels. For freshly frozen postmortem brain tissues, around 20 mg tissue was cut from the SN region and thoroughly homogenized in RIPA buffer supplemented with PI. The protein was extracted by incubating the samples in RIPA buffer overnight with constant rotation at 4°C. After gel electrophoresis, proteins from the gel were transferred to PVDF membrane by cold transfer electrophoresis. To detect α-SYN in human samples, only 15 μg protein per sample was used. The membranes were blocked by 5% fat-free milk prepared in Tris buffered saline (TBS)-Tween 20 (0.1%) for one hour and incubated overnight with primary antibodies prepared in blocking buffer as mentioned in the text. The membrane was washed 3 times with wash buffer (TBS-T) and incubated with secondary antibodies for 1 hour at RT. The protein bands were visualized by Enhanced Chemiluminescent (ECL) detection reagents (Super Signal West Pico Chemiluminescent Substrate, Thermo Scientific, 34077 or ECL Prime Western Blotting Detection Reagent, Amersham, 45-010-090).

### Total RNA isolation and cDNA synthesis

To measure expression from SH-SY5Y cells, total RNA isolation and consecutive cDNA synthesis were performed as described in our previous publication (Basu et al, 2017). The relative level of expression of each gene was normalized by β-actin gene expression. To measure expression from FANS-isolated neuronal nuclei, cDNA pre-amplification was performed prior to PCR amplification. Briefly, the collected nuclei were pelleted down immediately and lysed in 100 μL RA1 buffer from the Nucleo Spin RNA XS Kit (Takara, 740902.50). This kit is optimized for isolating RNA from as little as one cell. The RNA was eluted according to the protocol in 10 uL nuclease free water and equal amounts of RNAs from different samples were immediately subjected to amplification-based cDNA synthesis using the SMARTer PCR cDNA Synthesis Kit (Takara, 634925, 639207). We have standardized the amplification cycle to 15 for 2-10 ng of initial RNA template used for the reaction. The amplified cDNA was diluted 1:10 for gene-specific amplification. The purity of the collected NeuN-positive nuclei were tested by amplifying neuron-specific genes such as NeuN and synpatophysin. The samples were also checked for the presence of any astrocyte-specific gene expression such as GFAP. All the primers are listed in Supplementary Table 2.

### Statistical analyses

Statistical analyses were done using GraphPad Prism v.7 software (GraphPad Software Inc.). To compare between two groups, unpaired non-parametric student’s t-test was executed with Mann-Whitney post-hoc corrections. To measure correlation between two variables, Spearman correlation coefficient was calculated. Data are presented as mean ± SEM. Test values were considered significant if the p-value was ≤ 0.05.

## Acknowledgements

Authors gratefully acknowledge NIH Neurobiobank for providing all postmortem brain samples. Authors also thank Prof. Han-Seok Ko, Johns Hopkins University School of Medicine, for kindly providing all PD patient derived iPSC lines. Authors sincerely thank Prof. Viviane Labrie, Van Andel Research Institute, Prof. Rajiv Ratan, Burke Medical Research Institute and Prof. Flint M. Beal, Weill Cornell Medicine, for their review and critical scientific comments on the article. Financial support from NIH (1R01NS100919 awarded to YSK) is gratefully acknowledged. Authors also thank Mr. Andrew Knott for language editing, Mr. Levi Adams and Ms. Annlisa Simon for help with lentivirus generation, bacterial stock preparation, and plasmid DNA isolation and Ms. Anishaa Sivakumar for help with preparation of diagrammatic representation of figures.

## Author contributions

SGT and YSK conceptualized the study. SGT principally executed the work and developed all reagents. SGT and YSK wrote the manuscript. JK helped with iPSC culture and differentiation. SB helped with western blot analysis of the α-synuclein protein from postmortem brain tissues. SGT and MBF performed FANS-ChIP experiments and MBF helped with culturing of SH-SY5Y cells. EA and GJ initially helped with cloning of SunTag vectors. YSK supervised all the work.

## The Paper Explained

### PROBLEM

High levels of α-synuclein protein and its aggregation in the substantia nigra (SN) pars compacta of midbrain region is considered as the main culprit for degeneration of dopamine producing neurons in Parkinson’s disease. The main problem remains the lack of understanding about the molecular mechanisms how this protein gets expressed more in PD patients and how that initiates misfolding which leads to neurodegeneration. Several hypotheses have been put forward to understand α-synuclein’s aggregation behavior. However, there remain caveats in our understanding about de-regulated expression of this protein. Epigenetic regulations are one of the key mechanisms which regulate gene expression. However, how epigenetic factors underlying fine-tuning of α-synuclein expression get deregulated in the disease remains to be elucidated.

### RESULT

We observed the transcription-promoting histone post-translational modification (PTM), H3K4me3, is exclusively enriched at the α-synuclein gene promoter in postmortem brain samples of PD patients. The high levels of H3K4me3 at the promoter was positively correlated with higher α-synuclein protein levels in the SN of the brain. We have developed novel CRISPR/dCs9-based SunTag-JARID1A system which effectively reduced H3K4me3 enrichment from the α-synuclein gene promoter and concomitantly decreased the protein levels both in neuronal SHSY5Y cells as well as in dopaminergic neurons derived from PD patients’ iPSCs. These results implicate the importance of H3K4me3 in regulation of α-synuclein in PD.

### IMPACT

Epigenetic factors confer the first line of regulation for gene expression. Histone PTMs are the most important regulators for tissue-specific gene expression. We identified a histone PTM, H3K4m3, gets deregulated in PD which in turn perturbs α-synuclein expression in the patients. Genomic locus-specific editing of this epigenetic mark in cultured human neurons from patients significantly reduced α-synuclein levels. The impact of this novel approach indicates that H3K4me3 may serve as a molecular target to slow down synucleinopathy-mediated dopaminergic neuronal degeneration in PD.

## Data availability

The novel plasmids generated in the study will soon be deposited to Addgene. Any cell lines used in the study will be available from the authors upon request.

