## Supplemental materials for "Targeted attenuation of elevated histone marks at *SNCA* alleviates α-synuclein in Parkinson’s disease"

**List of Extended View Figures and tables**

Fig EV1. Distribution of histone PTM peaks in the regulatory region of human *SNCA*.

Fig EV2. H3K27ac and H3K27me3 did not show appreciable difference in enrichment at the *SNCA* promoter between control and PD brains.

Fig EV3. Enhancer-associated histone mark H3K27ac in *SNCA*-intron 4 not significantly different between control and PD.

Fig EV4. dCas9-5xGCN4 and scFV-sfGFP-JARID1A form a stable complex.

Fig EV5. Establishment of SH-SY5Y cells stably expressing dCas9-5xGCN4.

Fig EV6. dCas9-5xGCN4 was precisely recruited at the *SNCA* promoter as directed by sgRNAs.

Fig EV7. Relative efficiency of sgRNAs in reducing H3K4me3 from the *SNCA* promoter.

Fig EV8. Individual components of the CRISPR/dCas9 SunTag-JARID1A system do not affect α-synuclein or global H3K4me3 levels.

Fig EV9. Immunostaining of differentiated sPD iPSCs demonstrates successful differentiation to dopaminergic neurons.

Fig EV10. Sequence of scFV-sfGFP-JARID1A construct in pLvx vector.

Table EV1. Details of postmortem brain tissue samples used in the study.

Table EV2. List of primers used in the study.

Table EV3. List of short guide RNAs used in the study.

**Extended view Figure 1. Distribution of histone PTM peaks in the regulatory region of human *SNCA*.**

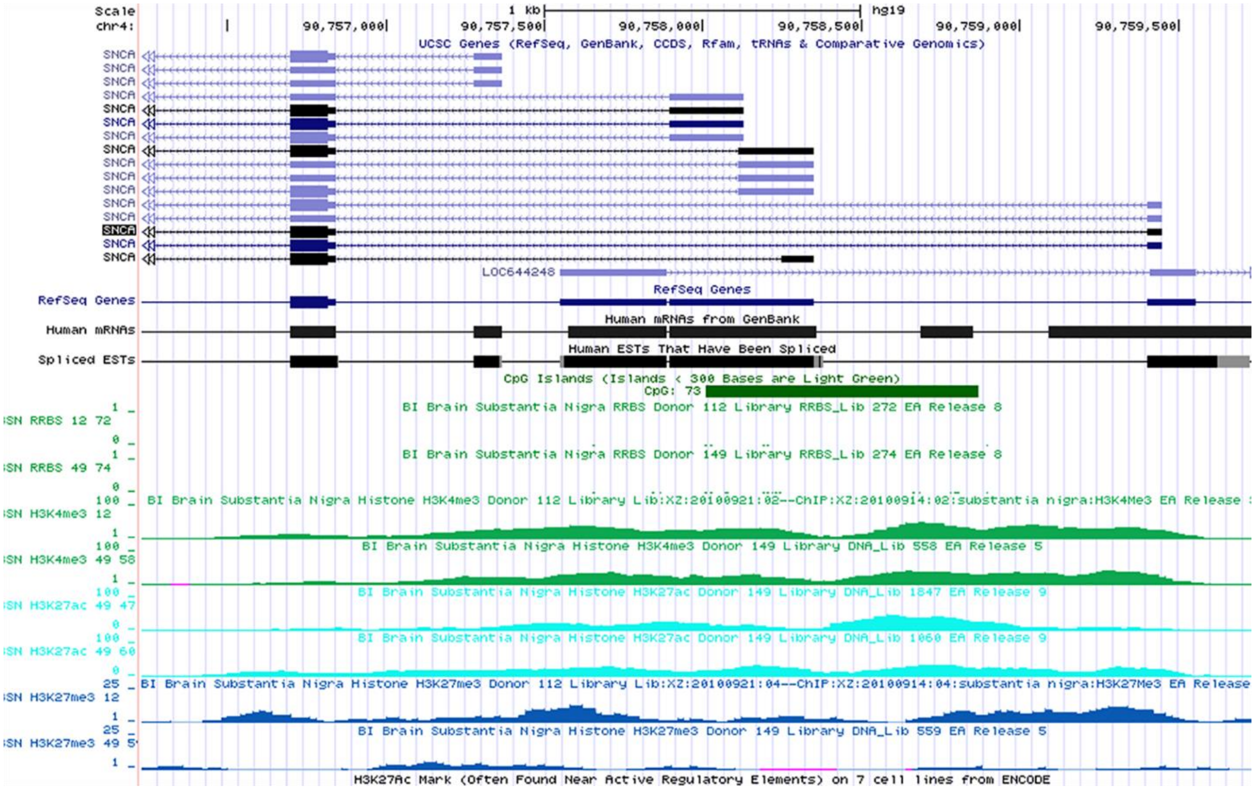


Screenshot from Roadmap Epigenomics Database depicts the distribution of histone PTMs (H3K4me3, H3K27me3, and H3K27ac) at the upstream regulatory region (promoter, its upstream regions, and intron 1) of *SNCA* on chromosome 4 from SN tissues of two healthy adult postmortem brain samples. The green horizontal bar represents the position of the CpG island at the *SNCA* promoter region as per the database (RRBS; reduced representation of bisulphite sequencing). Different transcripts (both coding and non-coding) of *SNCA* with their different TSS are shown by blue horizontal lines where boxes represent exons. The scale for the location of the genomic region from hg19 contig is also shown at the top. The screenshot only displays the genomic region of chromosome 4 from 90,756,000 to 90,760,000 bp. Arrows represent the direction of the gene.

**Extended view Figure 2. H3K27ac and H3K27me3 did not show appreciable difference in enrichment at the *SNCA* promoter between control and PD brains.**

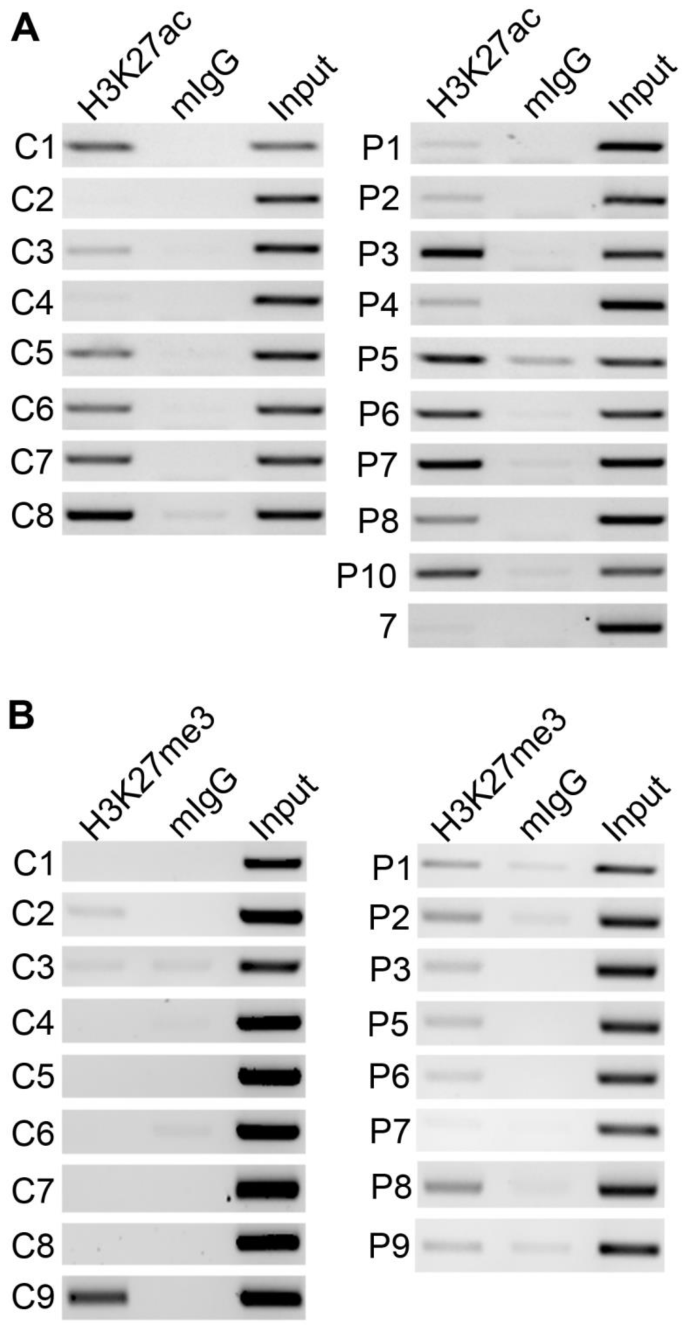


(**A**) ChIP gel images show relative enrichment by H3K27ac at the *SNCA* intron1 from 8 controls and 10 PD brains. No significant difference was observed (Extended view Figure 1D). (**B**) ChIP gel images demonstrating relative enrichment by H3K27me3 in 9 control and 8 PD subjects. The statistical difference between control and PD is shown in Figure 1C. For both H3K27ac and H3K27me3, PCR was performed for the same region as for H3k4me3. PCR amplified a 188-bp region from intron 1 of *SNCA* where these marks were found to be enriched in the ENCODE database.

**Extended view Figure 3. Enhancer-associated histone mark H3K27ac in *SNCA*-intron 4 not significantly different between control and PD.**

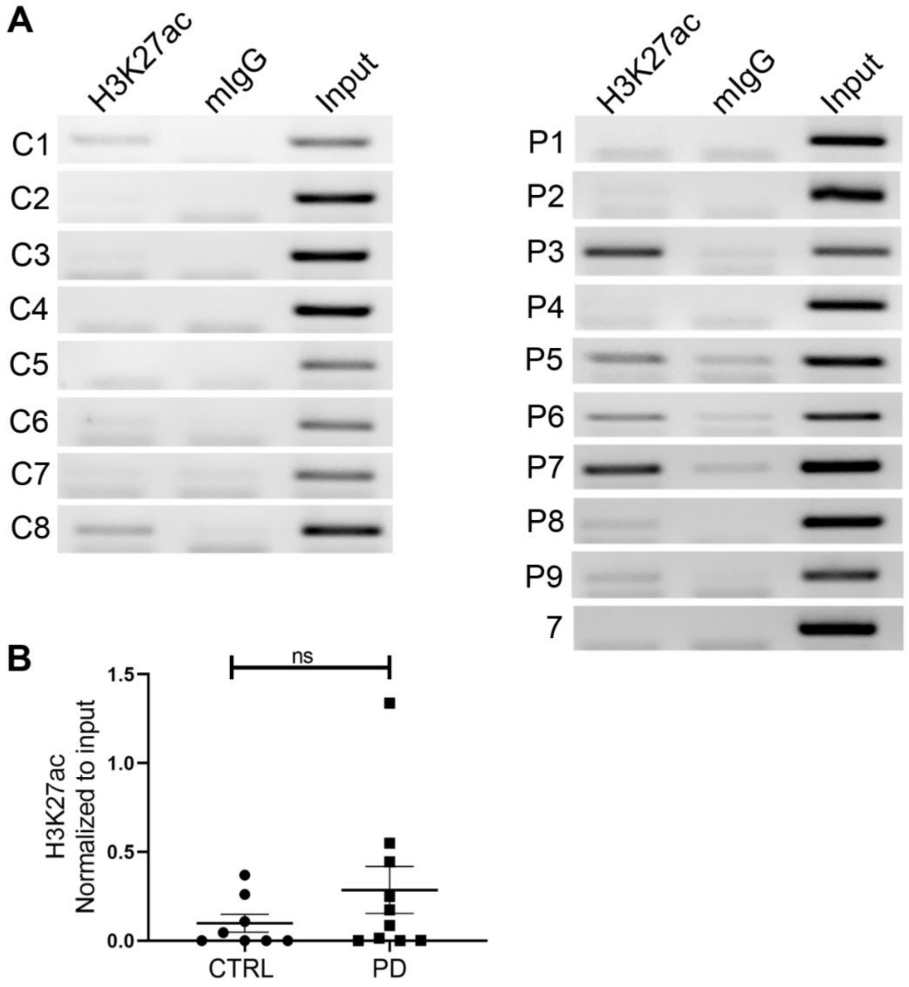


(**A**) ChIP gel images showing relative enrichment of H3K27ac in the intron 4 region of *SNCA* in control (n=8) and PD subjects (n=10). The PCR amplified 155-bp region in intron 4 corresponds to the H3K27ac peak in the ENCODE database. (**B**) Mean of the normalized intensities of relative H3K4me3 enrichment between control and PD were evaluated. No significant difference was observed between the groups. All data are presented as mean ± SEM. ns = non-significant. Data were analyzed using non-parametric t-test followed by Mann-Whitney post-hoc corrections. Two-tailed p-values were calculated.

**Extended view Figure 4. dCas9-5xGCN4 and scFV-sfGFP-JARID1A form a stable complex.**

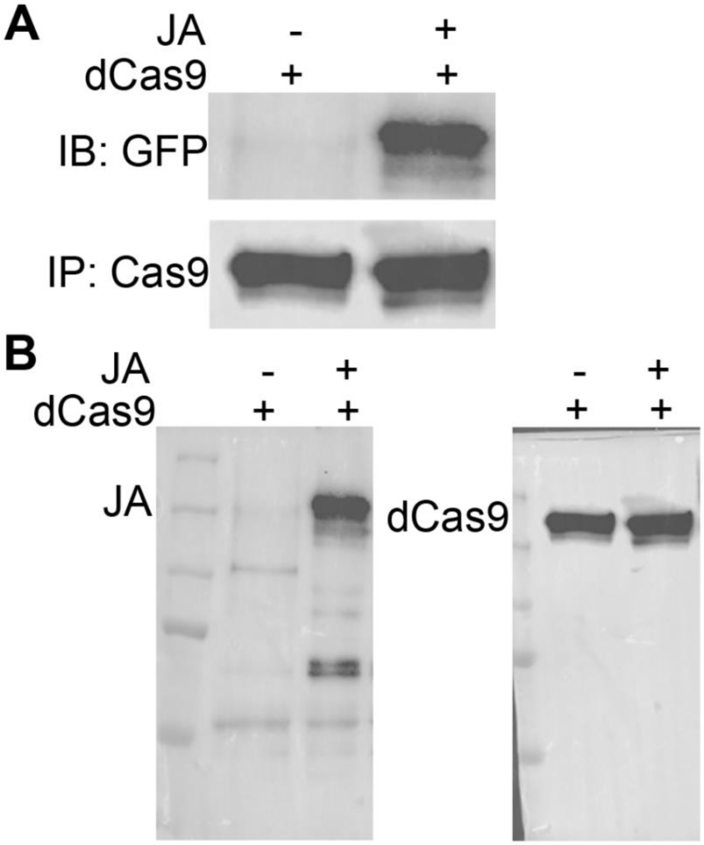


(**A**) dCas9-5xGCN4 plasmid was overexpressed in HEK293T cells with or without scFV-sfGFP-JARID1A. Total protein was isolated and immunoprecipitated with anti-cas9 antibody and immunoblotted against anti-GFP antibody. Cas9 antibody could pull down scFV-sfGFP-JARID1A (JA) (~168 kda) from HEK293T cells expressing both the constructs, whereas it failed to pull down any protein in other conditions. (**B**) Full western gel images of immunoblots used in A.

**Extended view Figure 5. Establishment of SH-SY5Y cells stably expressing dCas9-5xGCN4.**

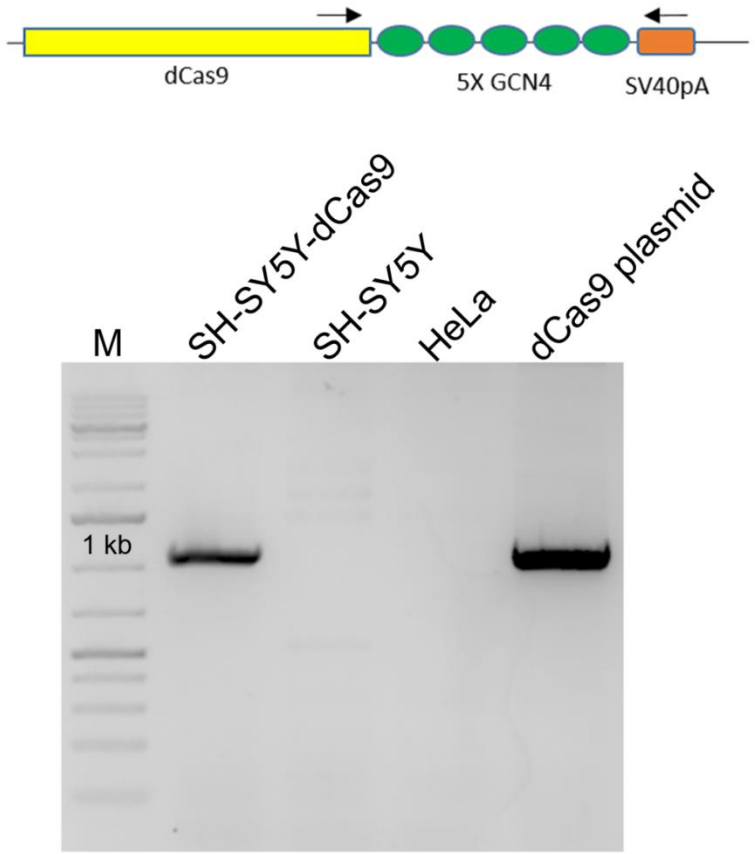


SH-SY5Y cells expressing dCas9-5xGCN4 were created by blasticidin selection. To ensure these cells expressed dCas9 and 5xGCN4, we performed PCR with forward and reverse primers just outside of dCas9-5xGCN4. The proper size band of ~1.1 kb was amplified from the genomic DNA isolated from stable SH-SY5Y cells expressing dCas9-5xGCN4. Two negative control cell lines, wild-type SH-SY5Y and HeLa cells, were used. The original dCas9-5xGCN4 plasmid was used as a positive control.

**Extended view Figure 6. dCas9-5xGCN4 was precisely recruited at the *SNCA* promoter as directed by sgRNAs.**

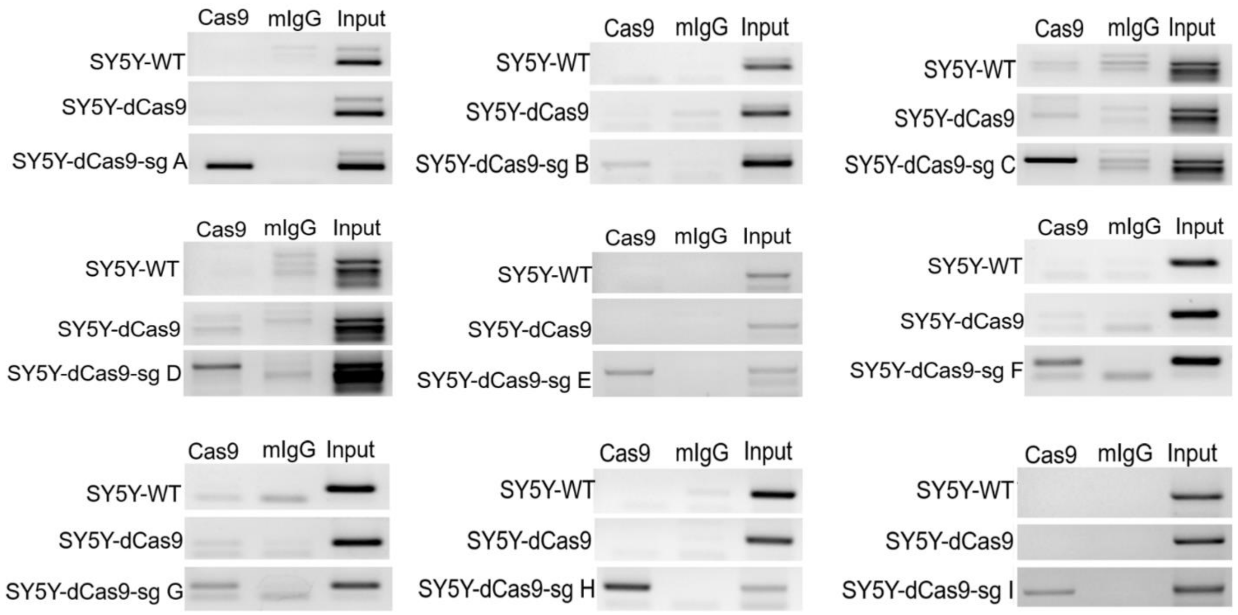


Each SH-SY5Y cell line stably expressing dCas9-5xGCN4 together with each sgRNA (a-i) was subjected to chromatin IP with cas9 antibody. Enrichment of dCas9-5xGCN4 was evaluated by PCR amplification using primer set encompassing the sgRNA binding sites. All the guide RNA sequences, corresponding primer sequences, and product sizes are listed in Extended view Table 3. Wild-type SH-SY5Y cells and SH-SY5Y-dCas9-5xGCN4 were checked in parallel as negative controls. All nine sgRNAs successfully recruited dCas9-5xGCN4 at the site.

**Extended view Figure 7. Relative efficiency of sgRNAs in reducing H3K4me3 from the *SNCA* promoter.**

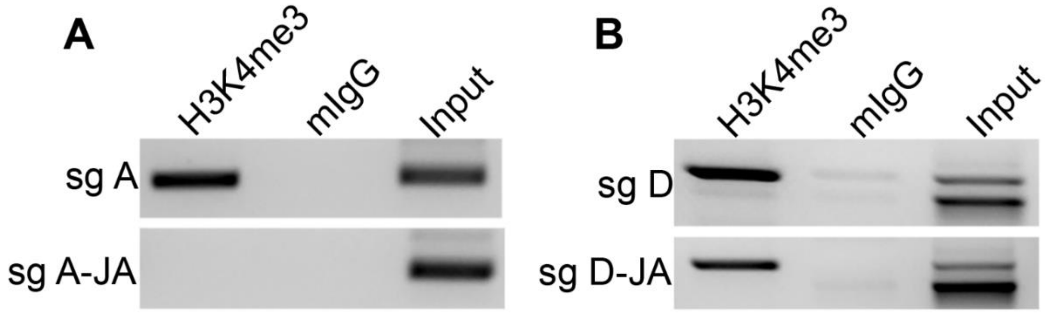


**(A, B)** ChIP gel images showing the enrichment of H3K4me3 between sgA and sgD. The efficiency of H3K4me3 reduction was compared in SH-SY5Y cells stably expressing sgA-dCas9-5xGCN4 (**A**) and sgD-dCas9-5xGCN4 (**B**) in presence of scFV-sfGFP-JARID1A. Results show sgA completely removed H3K4me3, while sgD showed partial reduction. Mouse IgG (mIgG) was used as an antibody control and 1% input was used as control for total chromatin.

**Extended view Figure 8. Individual components of the CRISPR/dCas9 SunTag-JARID1A system do not affect α-synuclein or global H3K4me3 levels.**

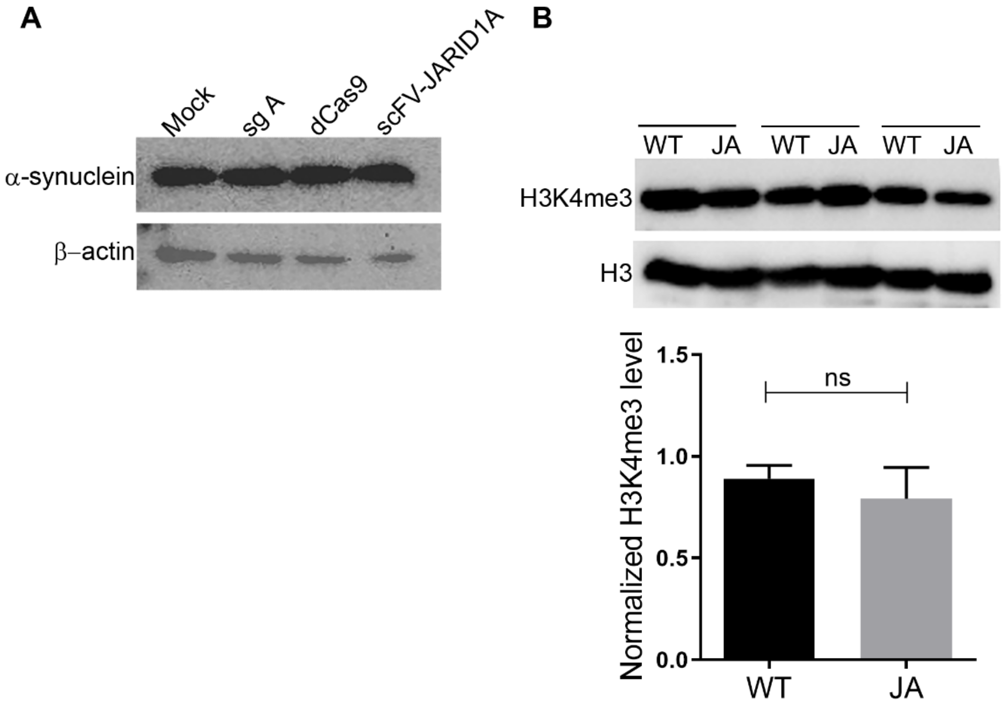


(**A**) The three components of the CRISPR/dCas9 SunTag-JARID1A system were individually overexpressed in SH-SY5Y cells. No difference in α-synuclein expression levels was observed in western blots. β-actin was used as an internal control. (**B**) SH-SY5Y cell line stably expressing sgA-dCas9-JARID1A (JA), and wild-type cells (WT) were compared for global H3K4me3 levels. No significant difference in global H3K4me3 levels was observed. Total H3 level was used as an endogenous control. Three independent repeats were performed (n=3). Data are presented as mean ± SEM. ns, no significant difference in the mean. Data were analyzed using non-parametric t-test followed by Mann-Whitney post-hoc corrections. Two-tailed p-values was calculated.

**Extended view Figure 9. Immunostaining of differentiated sPD iPSCs demonstrates successful differentiation to dopaminergic neurons.**

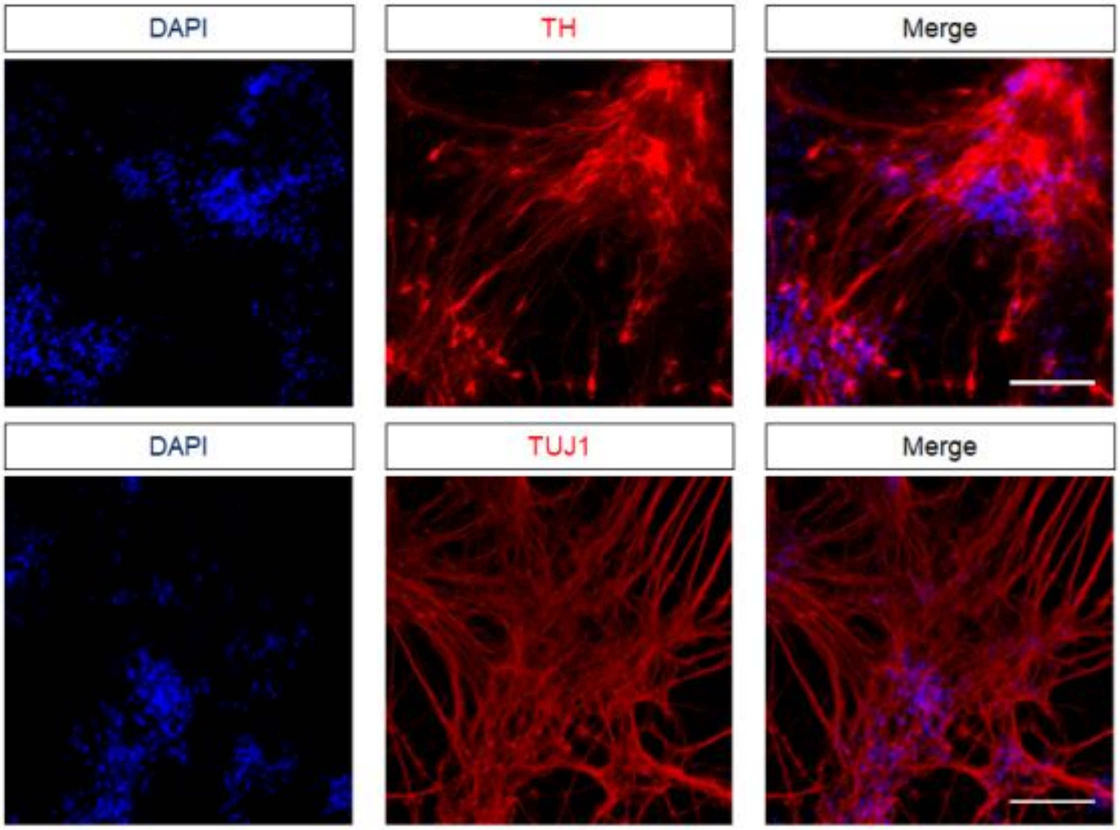


\

sPD1-1 line was differentiated for 30 days and subjected to immunostaining for the neuronal marker β-III tubulin (TUJ1) and TH, a dopaminergic neuron marker. Majority of the cells expressed TUJ1 and TH, confirming adequate differentiation into dopaminergic neurons. Hoechst was used to stain the nuclei. Scale bar, 100 µm.

**Extended view Figure 10. Sequence of scFV-sfGFP-JARID1A construct in pLvx vector.**

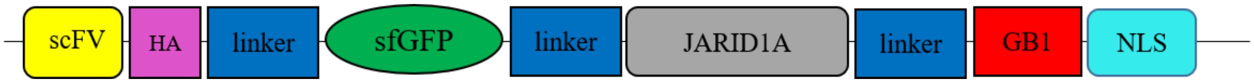


ATGGGCCCCGACATCGTGATGACCCAGAGCCCCAGCAGCCTGAGCGCCAGCGTGGGCGACCGCGTGACCATCACCTGCCGCAGCAGCACCGGCGCCG

TGACCACCAGCAACTACGCCAGCTGGGTGCAGGAGAAGCCCGGCAAGCTGTTCAAGGGCCTGATCGGCGGCACCAACAACCGCGCCCCCGGCGTGCC

CAGCCGCTTCAGCGGCAGCCTGATCGGCGACAAGGCCACCCTGACCATCAGCAGCCTGCAGCCCGAGGACTTCGCCACCTACTTCTGCGCCCTGTGG

TACAGCAACCACTGGGTGTTCGGCCAGGGCACCAAGGTGGAGCTGAAGCGCGGCGGCGGCGGCAGCGGCGGCGGCGGCAGCGGCGGCGGCGGCAGCA

GCGGCGGCGGCAGCGAGGTGAAGCTGCTGGAGAGCGGCGGCGGCCTGGTGCAGCCCGGCGGCAGCCTGAAGCTGAGCTGCGCCGTGAGCGGCTTCAG

CCTGACCGACTACGGCGTGAACTGGGTGCGCCAGGCCCCCGGCCGCGGCCTGGAGTGGATCGGCGTGATCTGGGGCGACGGCATCACCGACTACAAC

AGCGCCCTGAAGGACCGCTTCATCATCAGCAAGGACAACGGCAAGAACACCGTGTACCTGCAGATGAGCAAGGTGCGCAGCGACGACACCGCCCTGT

ACTACTGCGTGACCGGCCTGTTCGACTACTGGGGCCAGGGCACCCTGGTGACCGTGAGCAGCTACCCATACGATGTTCCAGATTACGCTGGTGGAGG

CGGAGGTTCTGGGGGAGGAGGTAGTGGCGGTGGTGGTTCAGGAGGCGGCGGAAGCTTGGATCCAGGTGGAGGTGGAAGCGGTAGCAAAGGAGAAGAA

CTTTTCACTGGAGTTGTCCCAATTCTTGTTGAATTAGATGGTGATGTTAATGGGCACAAATTTTCTGTCCGTGGAGAGGGTGAAGGTGATGCTACAA

ACGGAAAACTCACCCTTAAATTTATTTGCACTACTGGAAAACTACCTGTTCCGTGGCCAACACTTGTCACTACTCTGACCTATGGTGTTCAATGCTT

TTCCCGTTATCCGGATCACATGAAACGGCATGACTTTTTCAAGAGTGCCATGCCCGAAGGTTATGTACAGGAACGCACTATATCTTTCAAAGATGAC

GGGACCTACAAGACGCGTGCTGAAGTCAAGTTTGAAGGTGATACCCTTGTTAATCGTATCGAGTTAAAGGGTATTGATTTTAAAGAAGATGGAAACA

TTCTTGGACACAAACTCGAGTACAACTTTAACTCACACAATGTATACATCACGGCAGACAAACAAAAGAATGGAATCAAAGCTAACTTCAAAATTCG

CCACAACGTTGAAGATGGTTCCGTTCAACTAGCAGACCATTATCAACAAAATACTCCAATTGGCGATGGCCCTGTCCTTTTACCAGACAACCATTAC

CTGTCGACACAATCTGTCCTTTCGAAAGATCCCAACGAAAAGCGTGACCACATGGTCCTTCTTGAGTTTGTAACTGCTGCTGGGATTACACATGGCA

TGGATGAGCTCTACAAAGGTGGAGGTCGGACCGGTGGCGGTGGCGGAGGGATGGCGGGCGTGGGGCCGGGGGGCTACGCGGCGGAGTTCGTGCCACC

GCCAGAGTGCCCCGTCTTTGAGCCGAGTTGGGAGGAGTTCACAGATCCGCTCAGCTTTATCGGCCGCATCCGGCCTTTGGCGGAGAAAACCGGCATC

TGCAAAATTCGGCCGCCCAAGGACTGGCAGCCTCCATTTGCCTGTGAAGTAAAAAGCTTTCGTTTCACTCCAAGAGTCCAGCGCCTGAATGAACTTG

AGGCAATGACCAGAGTGAGATTGGATTTCTTGGATCAACTAGCAAAATTTTGGGAACTTCAAGGATCTACTCTGAAGATCCCTGTGGTAGAGAGAAA

AATCCTGGATCTGTATGCTTTGAGCAAGATTGTTGCCAGCAAAGGAGGTTTTGAAATGGTCACCAAAGAGAAGAAATGGTCTAAAGTGGGTAGTCGC

TTGGGATATCTGCCAGGAAAAGGAACTGGGTCTCTTTTGAAGTCACATTATGAAAGAATTCTCTACCCATATGAGCTTTTCCAGTCTGGTGTGAGCC

TTATGGGTGTGCAGATGCCTAATTTAGATCTTAAAGAAAAAGTGGAGCCTGAGGTTCTCAGCACTGATACCCAAACTTCCCCAGAGCCAGGCACAAG

GATGAACATTCTGCCGAAGAGAACAAGACGTGTGAAGACTCAGTCAGAATCTGGAGATGTGAGTAGAAACACGGAACTGAAGAAACTTCAGATTTTT

GGGGCTGGGCCCAAGGTTGTGGGCTTGGCAATGGGAACAAAAGATAAAGAAGATGAGGTCACCCGAAGACGAAAAGTTACCAACAGGTCAGACGCAT

TTAACATGCAAATGAGACAACGGAAAGGCACTCTCTCTGTTAACTTTGTTGATCTCTATGTTTGTATGTTTTGTGGTCGGGGAAACAATGAAGATAA

ATTGCTTTTGTGTGATGGATGTGATGACAGCTATCATACATTTTGTCTAATTCCTCCACTACCTGATGTGCCCAAAGGAGACTGGAGGTGTCCTAAA

TGTGTCGCCGAGGAATGTAGCAAACCTCGAGAAGCCTTTGGATTTGAACAAGCTGTACGAGAGTATACACTTCAGAGCTTTGGAGAGATGGCAGATA

ATTTTAAGTCTGATTATTTTAATATGCCAGTCCATATGGTTCCCACAGAACTAGTAGAAAAGGAATTTTGGCGGCTGGTAAGCAGCATTGAAGAAGA

TGTTATTGTGGAATATGGAGCAGATATCTCCTCAAAAGACTTTGGAAGTGGATTTCCGGTGAAGGATGGGCGGAGAAAGATTCTGCCAGAAGAAGAG

GAATATGCACTTTCTGGTTGGAATTTGAATAACATGCCTGTCCTGGAACAGTCTGTTCTTGCACATATTAATGTGGACATCTCTGGTATGAAAGTGC

CGTGGCTCTATGTGGGAATGTGCTTCTCTTCTTTTTGCTGGCACATTGAGGATCACTGGAGTTATTCCATCAACTACTTGCACTGGGGGGAGCCAAA

GACATGGTATGGTGTGCCATCTCATGCTGCAGAGCAACTGGAGGAGGTGATGAGAGAGCTGGCCCCCGAGTTATTTGAATCCCAGCCTGATCTTCTG

CATCAGTTAGTTACCATCATGAACCCCAACGTGCTAATGGAGCATGGTGTGCCTGTGTACAGGACCAATCAGTGTGCTGGCGAGTTTGTTGTGACAT

TTCCTCGTGCCTATCACTCTGGATTTAACCAGGGCTACAACTTTGCTGAAGCTGTGAACTTCTGTACTGCTGACTGGTTGCCCATTGGACGTCAATG

TGTAAATCATTACCGACGCCTAAGGCGCCACTGTGTCTTTTCACACGAGGAACTAATTTTCAAGATGGCAGCAGATCCAGAATGCTTAGATGTGGGG

CTGGCTGCCATGGTCTGCAAAGAATTGACTCTCATGACTGAAGAAGAAACACGATTAAGAGAGTCTGTTGTACAGATGGGTGTCCTGATGTCAGAAG

AAGAAGTGTTTGAACTTGTTCCTGATGATGAGCGGCAGTGTTCAGCATGCAGAACCACATGTTTTCTCTCTGCTCTCACATGTTCCTGTAATCCTGA

GCGGCTTGTATGTCTCTACCATCCAACTGATCTGTGCCCCTGCCCCATGCAGAAGAAATGTCTTAGATATCGCTACCCATTAGAAGACCTCCCTTCT

CTGCTATATGGTGTAAAAGTCAGGGCACAGTCCTATGACACTTGGGTCAGTCGTGTTACAGAAGCATTGTCTGCTAACTTCAACCACAAAAAAGATT

TGATTGAATTGCGAGTAATGCTGGAAGATGCTGAGGATAGGAAATACCCAGAGAATGATCTCTTTCGAAAACTCAGGGATGCTGTAAAAGAAGCTGA

GACCTGTGCTTCTGTGGGTAGTGGAGGAGGATCTCGGACCGAAGAGTACAAGCTTATCCTGAACGGTAAAACCCTGAAAGGTGAAACCACCACCGAA

GCTGTTGACGCTGCTACCGCGGAAAAAGTTTTCAAACAGTACGCTAACGACAACGGTGTTGACGGTGAATGGACCTACGACGACGCTACCAAAACCT

TCACGGTAACCGAAGGTGGTGGTAGCGGTGGTGGTACTAGTCCAAAAACAAGGAGGAGACCGCGAAGATCACAACGGAAAAGGCCGCCTACGCCATG

GCCGTAA

Organization of the scFV-sfGFP-JARID1A vector shown in Figure 4. The sequence of the entire insert cloned in a pLvx-DsRed vector is shown. The sequence of each component of the vector is color-coded both in the diagram and in the sequence.

**Extended view Table 1. Details of post-mortem brain tissue samples used in the study.**

| Source | Sample Number | Diagnosis | Age/Sex | Brain Region | PMI (h) |
|---|---|---|---|---|---|
| McLean Hospital, Harvard Medical School | 1 | Parkinson's Disease | 70/M | SN | 26.36 |
| McLean Hospital, Harvard Medical School | 2 | Parkinson's Disease | 79/M | SN | 10.1 |
| McLean Hospital, Harvard Medical School | 3 | Parkinson's Disease | 89/M | SN | 19.92 |
| McLean Hospital, Harvard Medical School | 4 | Parkinson's Disease | 85/M | SN | 24.75 |
| McLean Hospital, Harvard Medical School | 5 | Parkinson's Disease | 87/M | SN | 24.72 |
| McLean Hospital, Harvard Medical School | 6 | Parkinson's Disease | 74/M | SN | 26.98 |
| McLean Hospital, Harvard Medical School | 7 | Parkinson's Disease | 89/M | SN | 29 |
| McLean Hospital, Harvard Medical School | 8 | Parkinson's Disease | 74/M | SN | 35.42 |
| McLean Hospital, Harvard Medical School | 9 | Parkinson's Disease | 77/M | SN | 6.62 |
| McLean Hospital, Harvard Medical School | 10 | Parkinson's Disease | 71/M | SN | 15.67 |
| Human Brain and Spinal Fluid Resource Centre, Brain bank, UCLA | P1 | Parkinson's Disease | 73/M | SN | 15 |
| Human Brain and Spinal Fluid Resource Centre, Brain bank, UCLA | P2 | Parkinson's Disease | 79/M | SN | 12.3 |
| Human Brain and Spinal Fluid Resource Centre, Brain bank, UCLA | P3 | Parkinson's Disease | 82/M | SN | 13.0 |
| Human Brain and Spinal Fluid Resource Centre, Brain bank, UCLA | P4 | Parkinson's Disease | 83/M | SN | 6.7 |
| Human Brain and Spinal Fluid Resource Centre, Brain bank, UCLA | P5 | Parkinson's Disease | 78/M | SN | 13 |
| Human Brain and Spinal Fluid Resource Centre, Brain bank, UCLA | P6 | Parkinson's Disease | 87/M | SN | 11.2 |
| Human Brain and Spinal Fluid Resource Centre, Brain bank, UCLA | P7 | Parkinson's Disease | 81/M | SN | 9.7 |

| | | | | | |
|---|---|---|---|---|---|
| Human Brain and Spinal Fluid Resource Centre, Brain bank, UCLA | P8 | Parkinson’s Disease | 75/M | SN | 13.8 |
| Human Brain and Spinal Fluid Resource Centre, Brain bank, UCLA | P9 | Parkinson’s Disease | 83/M | SN | 16.3 |
| Brain Endowment Bank, University of Miami, Miller School of Medicine | C1 | Control | 89/M | SN | 20.5 |
| Brain Endowment Bank, University of Miami, Miller School of Medicine | C2 | Control | 87/M | SN | 10 |
| Brain Endowment Bank, University of Miami, Miller School of Medicine | C3 | Control | 84/M | SN | 27.5 |
| Brain Endowment Bank, University of Miami, Miller School of Medicine | C4 | Control | 54/M | SN | 26.5 |
| Brain Endowment Bank, University of Miami, Miller School of Medicine | C5 | Control | 76/M | SN | 30.25 |
| Brain Endowment Bank, University of Miami, Miller School of Medicine | C6 | Control | 70/M | SN | 27.2 |
| Brain Endowment Bank, University of Miami, Miller School of Medicine | C7 | Control | 76/M | SN | 27.75 |
| Brain Endowment Bank, University of Miami, Miller School of Medicine | C8 | Control | 55/M | SN | 26.2 |
| Brain Endowment Bank, University of Miami, Miller School of Medicine | C9 | Control | 75/M | SN | 14.16 |

**Extended view Table 2. List of primers used in the study.**

| Target | Experiment | 5'-3' | Product size (bp) |
|---|---|---|---|
| SNCA (promoter/intron1) | ChIP | F: TCCCCGGGAAACGCGAGGAT<br>R: CCCCGCGCCAGCACTTGTTA | 188 |
| SNCA (intron4) | ChIP | F:TGCCTTTGCATCAGATAATGGC<br>R:ATGATGAGCAGGCAGTCCG | 155 |
| α-synuclein | RNA | F:CACCATGGATGTATTCATGAA<br>R:AAAGATATTTCTTAGGCTTCAG | 437 |
| NeuN | RNA | F:CCCTTGCCGCTGGCTC<br>R:GCGACCACAGGAAGACTGTTA | 168 |
| Synpatophysin | RNA | F:TGCCAACAAGACCGAGAGTG<br>R:CAGAGCCCCCATGGAGTAGA | 196 |
| GFAP | RNA | F:GCACGCAGTATGAGGCAATG<br>R:TAGTCGTTGGCTTCGTGCTT | 139 |
| GAPDH | RNA | F:TTGCCATCAATGACCCCTTCA<br>R:TCCAAAATCAAGTGGGGCG | 153 |
| β-actin | RNA | F:GGAGTCCTGTGGCATCCACG<br>R:CTAGAAGCATTTGCGGTGGA | 322 |

**Extended View Table 3. List of short guide RNAs used in the study.**

| Guide RNA | Sequence (5'-3') | Validation primer sequence (5'-3') | PCR product size (bp) |
|---|---|---|---|
| sgA | AAGCAAAGGCTTTCTGCTAG (sense) | F: AGCGCAAGAATCAGACAAAGC<br>R:AGAATGGAGAAGCAAGCTCCTC | 179 |
| sgB | TCCGGTAGGCTAAATCACGC (sense) | F:AGTCAGAAAGGTGAGTGGTGTGTAG<br>R:AGCGGTCCTAAGGCTTTTCGCTCTAG | 164 |
| sgC | CCGCTTGTTTTAGACGGCTG (sense) | F:AGTCAGAAAGGTGAGTGGTGTGTAG<br>R:AGCCGGAAAGGGTCCTGAGGG | 357 |
| sgD | TGGGAAAATCAGCGTCTGGC (sense) | F:AGTCAGAAAGGTGAGTGGTGTGTAG<br>R:AGCCGGAAAGGGTCCTGAGGG | 357 |
| sgE | GCCGCGCAAGGCGGGAAAGT (antisense) | F:AGCAGCTCCCCAAGGGATAGGCTC<br>R:CACGCACCTCACTTCCGCGT | 521 |
| sgF | AAGGGCAGACCAATAGTTCA (sense) | F:CAAGGTCTCAAAGCCAGACAGCA<br>R:CTTGAACTATTGGTCTGCCCTTTGGA | 138 |
| sgG | CAAGTCCAACCTTCTTGCTC (sense) | F:CAAGGTCTCAAAGCCAGACAGCA<br>R:CTTGAACTATTGGTCTGCCCTTTGGA | 138 |
| sgH | GCGACTCTGACGAGGGGTAG (sense) | F:AGCAGCTCCCCAAGGGATAGGCTC<br>R:TGGAGATCGGGAGCGGTTGGGCTAG | 189 |
| sgI | ACTTTAAAACCACAAGGAAC (antisense) | F:AGCAGCTCCCCAAGGGATAGGCTC<br>R:AGAATGGAGAAGCAAGCTCCTC | 521 |
